# Accurate chromosome segregation is more dependent on kinetochore-bound Stu2 in meiosis than mitosis

**DOI:** 10.64898/2026.09.06.749677

**Authors:** Batula Robow, Soni Lacefield

**Author notes:** Corresponding author 66 College St. HB7200 Hanover, NH 03755.

## Abstract

Accurate chromosome segregation requires bipolar kinetochore-microtubule attachments. Error correction mechanisms promote cycles of kinetochore-microtubule release and reattachment until bipolar attachments are established. Although conserved error correction pathways operate in mitosis and meiosis, whether they have distinct requirements during meiosis remains unclear. Here, we investigate the role of Stu2/XMAP215/chTOG in kinetochore-microtubule error correction and compare its function with that of Ipl1/Aurora B during budding yeast mitosis and meiosis. Reducing kinetochore-bound Stu2 causes substantially more chromosome missegregation during meiosis than mitosis, revealing a heightened requirement for Stu2-mediated error correction. Whereas Ipl1-depleted cells fail to release initial improper kinetochore-microtubule attachments, kinetochore-defective *stu2* mutants undergo repeated error correction attempts yet missegregate chromosomes. Our findings reveal distinct but complementary functions for Ipl1 and Stu2. Ipl1 releases initial improper attachments, while kinetochore-bound Stu2 promotes the formation and stabilization of productive attachments and enables error correction attempts to establish biorientation. We propose that features of meiotic chromosome segregation increase the requirement for multiple error correction pathways.

## Introduction

Accurate chromosome segregation during mitosis and meiosis depends on the establishment of correct kinetochore-microtubule attachments, as well as the timely destabilization of erroneous attachments. Kinetochores initially associate with the lateral surface of spindle microtubules and are transported toward the spindle poles by motor proteins before converting to stable end-on attachments (Li et al., 2024). In mitosis, sister chromatid kinetochores establish bipolar attachments to microtubules from opposite spindle poles before anaphase onset. Meiosis imposes distinct constraints on this process. During meiosis I, sister kinetochores are co-oriented and function as a single microtubule-binding unit such that homologous chromosomes biorient. During meiosis II, sister kinetochores establish bipolar attachments to microtubules from opposite spindle poles, similar to mitosis. Erroneous attachments arise when sister kinetochores in mitosis or meiosis II, or homologous chromosome kinetochores in meiosis I, attach to microtubules emanating from the same spindle pole (Li et al., 2024; Cairo and Lacefield, 2020; Vukušić and Tolić, 2022). Conserved error correction pathways selectively destabilize erroneous attachments while simultaneously stabilizing bipolar, tension-bearing attachments to ensure faithful chromosome segregation. Careful comparison of error correction mechanisms during mitosis and meiosis revealed how conserved regulators are adapted to distinct kinetochore structures and chromosome segregation programs.

In both mitosis and meiosis, most chromosomes establish initial kinetochore-microtubule attachments to one pole during prometaphase, which are subsequently converted into bipolar, tension-bearing attachments (Li et al., 2024; Cairo and Lacefield, 2020; Vukušić and Tolić, 2022). Tension arises through the poleward pulling forces exerted by spindle microtubules, which are opposed by pericentromeric cohesion between sister chromatids in mitosis and meiosis II, or by crossover linkages together with arm cohesion between homologous chromosomes in meiosis I. Consequently, low tension attachments are repeatedly released and reformed until stable biorientation is achieved. Because error correction transiently generates unattached kinetochores, the spindle assembly checkpoint is activated, delaying anaphase onset and providing additional time for formation of productive attachments (McVey et al., 2021; Bhalla and Lacefield, 2026). Once bipolar, tension-bearing attachments are established, error correction is silenced, the spindle checkpoint is satisfied, and cells then proceed into anaphase.

Error correction is mediated by several conserved proteins including Ipl1/Aurora B kinase, Mps1 kinase, and the Stu2/XMAP215/chTOG microtubule polymerase (Li et al., 2024). Ipl1/Aurora B, as part of the chromosome passenger complex (CPC), is a key regulator of error correction that phosphorylates outer kinetochore proteins to destabilize improper kinetochore-microtubule interactions (McVey et al., 2021; DeLuca et al., 2006; Cheeseman et al., 2006; Dewar et al., 2004; Biggins et al., 1999; Tanaka et al., 2002). Previous studies provided evidence that Mps1 contributes to error correction, although this activity has been difficult to resolve from its well-established role in spindle checkpoint signaling (Santaguida et al., 2010; Pleuger and Westermann, 2026; Jones et al., 2005; Hewitt et al., 2010; Maciejowski et al., 2010; Jelluma et al., 2008; Maure et al., 2007; Benzi et al., 2020). More recently, a more direct role for Mps1 in error correction has been revealed, with Mps1 phosphorylating outer kinetochore proteins to weaken kinetochore-microtubule interactions that lack tension (Sarangapani et al., 2021; Maciejowski et al., 2017; Meyer et al., 2018; Leça et al., 2025). Ipl1/Aurora B and Mps1 act in complementary error correction pathways, and one cannot compensate for the loss of the other (Sarangapani et al., 2021). Finally, Stu2/XMAP215/chTOG is a conserved microtubule polymerase that can also promote microtubule destabilization and regulates plus-end dynamics (Li et al., 2024; Humphrey et al., 2018; Kosco et al., 2001; Gard and Kirschner, 1987; Brouhard et al., 2008; Cullen et al., 1999; Shirasu-Hiza et al., 2003; Podolski et al., 2014; Usui et al., 2003; Kitamura et al., 2010). Stu2/chTOG also localizes to kinetochores, where it promotes establishment of kinetochore biorientation (Miller et al., 2016; Zahm et al., 2021; Miller et al., 2019; Herman et al., 2020). *In vitro*, Stu2 ensures tension sensitivity to kinetochore-microtubule attachments, stabilizing tension-bearing attachments and destabilizing low tension attachment at disassembling microtubules (Miller et al., 2016). Moreover, combined perturbation of kinetochore-associated Stu2 and Ipl1 activity results in additive growth defects and elevated chromosome missegregation in budding yeast, consistent with these proteins acting through distinct but complementary mechanisms (Miller et al., 2019). Collectively, these findings support a model in which the three error correction mechanisms operate as independent pathways.

During meiosis, error correction mechanisms have evolved specialized functions. In budding yeast, homologous chromosome pairs frequently enter prometaphase attached to a single pole and must be actively reoriented to achieve bipolar attachment (Monje-Casas et al., 2007; Meyer et al., 2013). Ipl1 destabilizes kinetochore-microtubule attachments in prometaphase I, thereby enabling error correction and biorientation (Meyer et al., 2013). Mps1 is dispensable for this release but is essential for subsequent steps, including the clustering of kinetochores at the spindle pole body (SPB) during prophase I exit, formation of force-generating end-on attachments and efficient pole-to-pole chromosome movement (Meyer et al., 2018, 2013, 2021). Little is known about the meiotic functions of Stu2/XMAP215/chTOG in error correction of kinetochore-microtubule attachments. In fission yeast, the TOG family homolog Alp14, with its partner Alp7, helps capture kinetochores on chromosomes that become scattered during prophase I, allowing them to be retrieved to the spindle before meiosis I chromosome segregation (Kakui et al., 2013). In budding yeast meiosis, Stu2 has been shown to regulate microtubule dynamics, similar to its role in mitotic cells (Meyer et al., 2021).

Here, we use budding yeast to investigate the role of kinetochore-bound Stu2 in error correction across mitosis and meiosis and compare its function with that of Ipl1. We examine *stu2* mutant alleles that specifically disrupt its interaction with the outer kinetochore proteins in the Ndc80 complex while preserving its roles in regulating microtubule polymerization and dynamics (Zahm et al., 2021). These mutant alleles were initially thought to abolish Stu2 kinetochore binding, but recent studies suggest that additional, weaker interactions may provide alternative modes of kinetochore interaction (Abouelghar et al., 2025; Maitra et al., 2026). We found that the strong reduction of kinetochore-bound Stu2 causes substantially higher rates of chromosome missegregation in meiosis I and meiosis II than in mitosis, revealing an increased requirement for Stu2-mediated error correction during meiosis. Remarkably, despite repeated attempts at error correction and activation of the spindle checkpoint, *stu2* mutants frequently segregate chromosomes asymmetrically, with an excess of chromosomes partitioning toward either SPB. Together, our findings identify kinetochore-bound Stu2 as a critical component of meiotic error correction and reveal a distinct requirement for Stu2 in ensuring accurate chromosome segregation following the release of erroneous kinetochore-microtubule attachments.

## Results

### Loss of Ipl1 and kinetochore-bound Stu2 have opposing effects on the duration of metaphase I and metaphase II

Our previous work uncovered an unexpected difference in the requirement for Ipl1 in regulating cell-cycle timing during meiosis and mitosis. Depletion of Ipl1 accelerates anaphase onset in both meiosis I and meiosis II, but delays anaphase onset during mitosis (Cairo et al., 2020, 2023). Because Stu2 also regulates kinetochore-microtubule attachments, we asked whether loss of kinetochore-bound Stu2 similarly has distinct consequences for meiotic and mitotic progression. To this end, we analyzed two *stu2* mutant alleles in which Stu2 kinetochore-localization was disrupted while preserving its known function as a microtubule polymerase (Figure 1A). The *stu2^Δ855-888^*allele contains a C-terminal deletion that disrupts Stu2 binding to the Ndc80 kinetochore protein complex, whereas the *stu2^EEE^* allele contains three amino acid substitutions within this region that similarly disrupts the Stu2-Ndc80 interaction (Zahm et al., 2021).

**Figure 1.**
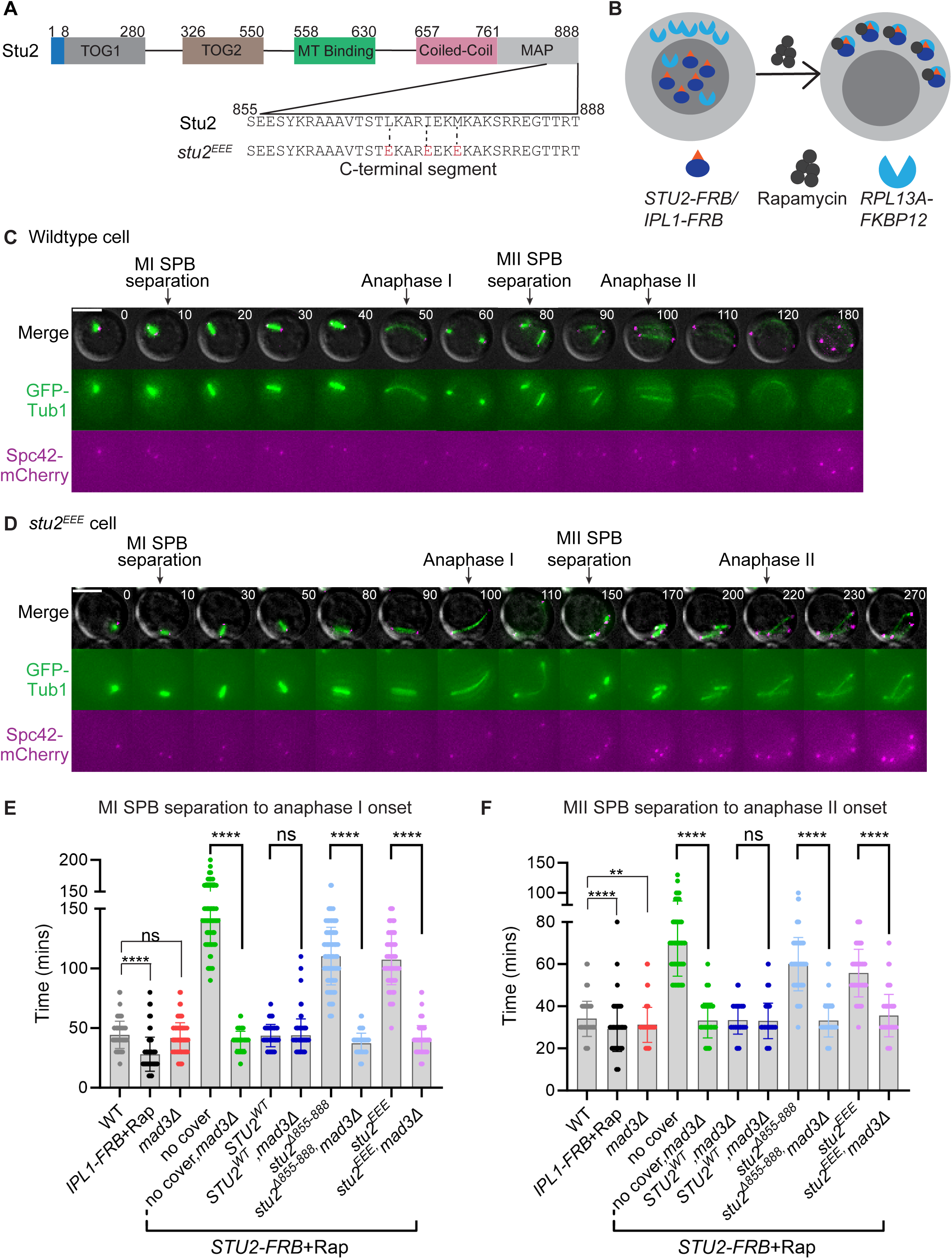
Stu2 depletion causes a spindle checkpoint-dependent delay in meiosis I and meiosis II. (A) Schematic representation of Stu2 protein functional domains showing the C-terminal segment that is needed for Stu2 kinetochore interaction. MT, microtubule. (B) Schematic of the Anchor Away technique to deplete either Stu2-FRB or Ipl1-FRB from the nucleus by addition of rapamycin (Rap) and coupling to a ribosomal receptor protein, Rpl13A-FKBP12 that shuttles from the nucleus into the cytosol. (C and D) Representative meiotic time-lapse images of a wildtype cell (C) or a cell with Stu2 nuclear depletion while expressing *stu2^EEE^* (D). Cells express Spc42-mCherry and GFP-Tub1. Scale bars, 5 μm. (E and F) Graph of the mean time (in minutes) from SPB separation in meiosis I to anaphase I onset (E) or from SPB re-duplication in meiosis II to anaphase II onset (F) in the indicated genotypes. Asterisks indicate a statistically significant difference between *stu2* mutant cells and *stu2 mad3Δ* mutant cells (****, P < 0.0001; **, P = 0.0019; ns, not significant; Mann-Whitney test). Error bars represent the standard deviation (SD); n ≥100 cells analyzed per genotype.

*STU2* and *IPL1* are essential genes, therefore, we employed the anchor-away system to acutely deplete Stu2 and Ipl1 from the nucleus at meiotic entry (Biggins et al., 1999; Chan and Botstein, 1993; Wang and Huffaker, 1997; Haruki et al., 2008)(Figure 1B). For this purpose, endogenous Stu2 and/or Ipl1 were C-terminally tagged with FRB, and the ribosomal protein Rpl13a tagged with FKBP12 in the same strain. Addition of rapamycin induces FRB-FKBP12 dimerization, rapidly sequestering FRB-tagged Stu2 and/or Ipl1 from the nucleus to the cytoplasm. In Stu2 anchor-away strains, *STU2* alleles (*STU2^WT^*, *stu2^Δ855-888^*, *stu2^EEE^*) were integrated at an ectopic locus and compared to a strain with no ectopic *STU2* allele (no cover). Cells with FRB-tagged Stu2 and Ipl1 grew comparably to wildtype cells in the absence of rapamycin, indicating that the tags did not impair protein function (Figure S1). As expected, strains with FRB-tagged Stu2 or Ipl1 exhibited a growth defect with addition of rapamycin, consistent with their efficient depletion from the nucleus. Importantly, expression of an ectopic copy of either wildtype or kinetochore-defective *stu2* alleles (*stu2^Δ855-888^*, *stu2^EEE^*) rescued this rapamycin-dependent growth defect to differing extents (Figure S1).

To determine whether loss of kinetochore-bound Stu2 impacts meiotic timing, we performed time-lapse microscopy and measured the interval between spindle pole body (SPB) separation (Spc42-mCherry) and anaphase spindle elongation (GFP-Tub1) during meiosis I and meiosis II (Figure 1C-F). As previously shown, Ipl1 depletion accelerated anaphase onset in both meiotic divisions compared with wildtype controls (Cairo et al., 2020). In striking contrast, depletion of Stu2 in cells expressing the kinetochore-defective *stu2* alleles substantially delayed anaphase onset. During meiosis I, anaphase onset occurred at 44 ± 11 minutes in wildtype cells, 142 ± 22 minutes in cells lacking a covering *STU2* allele (no cover), 110 ± 24 minutes in *stu2^Δ855-888^*, and 107 ± 21 minutes in *stu2^EEE^* cells. A similar delay was observed during meiosis II, with each mutant exhibiting delayed anaphase onset relative to wildtype controls (Figure 1C-F). Thus, unlike Ipl1 depletion, which accelerates meiotic progression, loss of kinetochore-bound Stu2 delays anaphase onset.

Activation of the spindle checkpoint delays anaphase in response to unattached kinetochores to provide additional time for proper kinetochore-microtubule attachments to form (Pleuger and Westermann, 2026). To determine whether the anaphase delay we observed in the absence of kinetochore-bound Stu2 requires spindle checkpoint activity, we monitored meiotic progression in cells lacking Mad3, a core component of the spindle checkpoint. Deletion of *MAD3* abolished the anaphase delay in the *stu2* mutants, demonstrating that the delay is spindle checkpoint dependent in both meiosis I and meiosis II (Figure 1E-F). Kinetochore-defective *stu2* alleles similarly cause a spindle checkpoint dependent delay during mitosis (Zahm et al., 2021). Together, these results indicate that kinetochore-bound Stu2 is required for the timely establishment of proper kinetochore-microtubule attachments during meiosis and suggest that loss of Stu2 at the kinetochore results in unattached kinetochores that activate the spindle checkpoint.

### Co-depletion of Ipl1 and Stu2 partially suppresses the spindle checkpoint-mediated anaphase delay caused by loss of kinetochore-bound Stu2

In contrast to Ipl1 depletion, which does not trigger a spindle checkpoint-dependent delay, loss of kinetochore-bound Stu2 activates the spindle checkpoint and delays anaphase onset (Miller et al., 2019; Zahm et al., 2021; Biggins and Murray, 2001)(Figure 1E-F). We therefore asked how simultaneous nuclear depletion of Ipl1 and kinetochore-bound Stu2 affects meiotic progression. Using the anchor-away system, we co-depleted endogenous Ipl1 and Stu2 in cells expressing kinetochore-defective *stu2* mutant alleles. Anaphase onset in both meiosis I and meiosis II in these cells occurred substantially earlier than following Stu2 depletion alone but remained delayed relative to wildtype cells. Mean anaphase I onset occurred at 79 ± 32minutes in the no-covering allele, 57 ± 22 minutes in *stu2^Δ855-888^*, and 55 ± 23 minutes in *stu2^EEE^* (Figure 2A-B). These results demonstrate that depletion of Ipl1 partially suppresses the delay caused by loss of kinetochore-bound Stu2.

**Figure 2.**
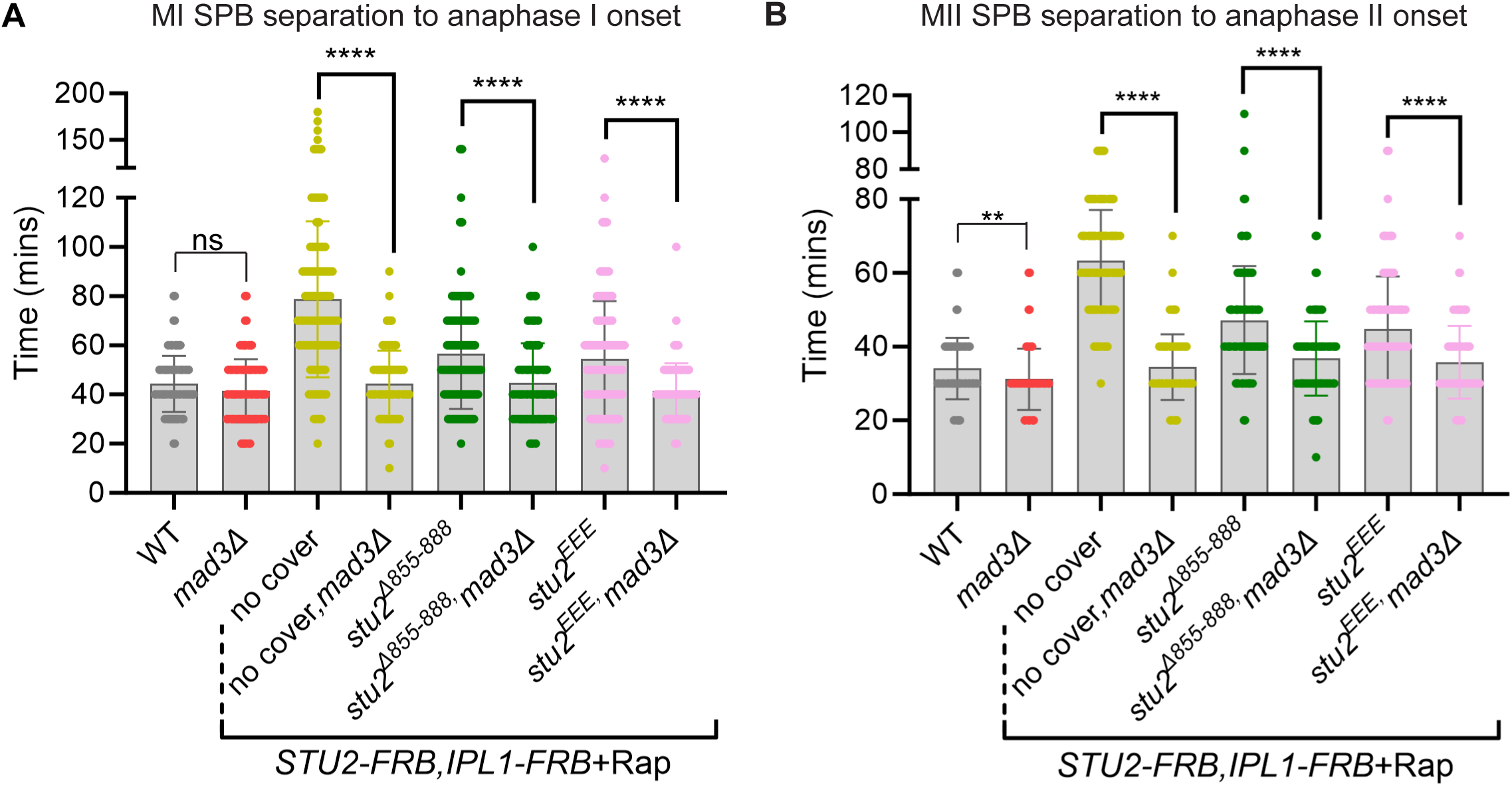
Ipl1 and Stu2 co-depletion delays anaphase onset, but not as severely as Stu2 depletion. (A and B) Graph of the mean time (in minutes) from SPB separation in meiosis I to anaphase I onset (A) or from SPB re-duplication in meiosis II to anaphase II onset (B) in the indicated genotypes. Asterisks indicate statistically significant differences between Stu2- and Ipl1-depleted cells with and without *mad3Δ* (****, P < 0.0001; **, P = 0.0019; ns, not significant; Mann-Whitney test). Error bars represent the standard deviation (SD); n ≥100 cells analyzed per genotype. Rap, Rapamycin.

We tested whether the residual delay in cells co-depleted of Ipl1 and Stu2 was spindle checkpoint dependent. Deletion of *MAD3* in cells co-depleted of Stu2 and Ipl1, including those expressing kinetochore-defective *stu2* alleles, restored anaphase I and anaphase II timing similar to wildtype levels, indicating that the residual delay is largely mediated by the spindle checkpoint (Figure 2A-B). Notably, the checkpoint-dependent delay was shorter following co-depletion of Ipl1 and Stu2 than in cells lacking kinetochore-bound Stu2 alone. This shorter delay is consistent with more rapid satisfaction of the spindle checkpoint, potentially through faster establishment of kinetochore-microtubule attachments. However, checkpoint satisfaction does not establish whether these attachments are correctly bioriented because they may be unable to release erroneous attachments.

We further investigated how co-depletion of Ipl1 and kinetochore-bound Stu2 affects mitotic progression (Figure 3A-D). In contrast to meiosis, Ipl1 depletion in mitosis anaphase onset independent of the spindle checkpoint (Cairo et al., 2020, 2023; Biggins and Murray, 2001) (Figure 3D). Loss of kinetochore-bound Stu2 causes a spindle checkpoint-dependent delay in anaphase onset (Zahm et al., 2021). With co-depletion of Ipl1 and Stu2, anaphase onset occurred with similar timing as cells depleted of Ipl1 alone and was substantially earlier than what we observed following Stu2 depletion alone (Figure 3A-D). These results suggest that depletion of Ipl1 rescues the spindle checkpoint-dependent delay caused by loss of kinetochore-bound Stu2 in mitosis.

**Figure 3.**
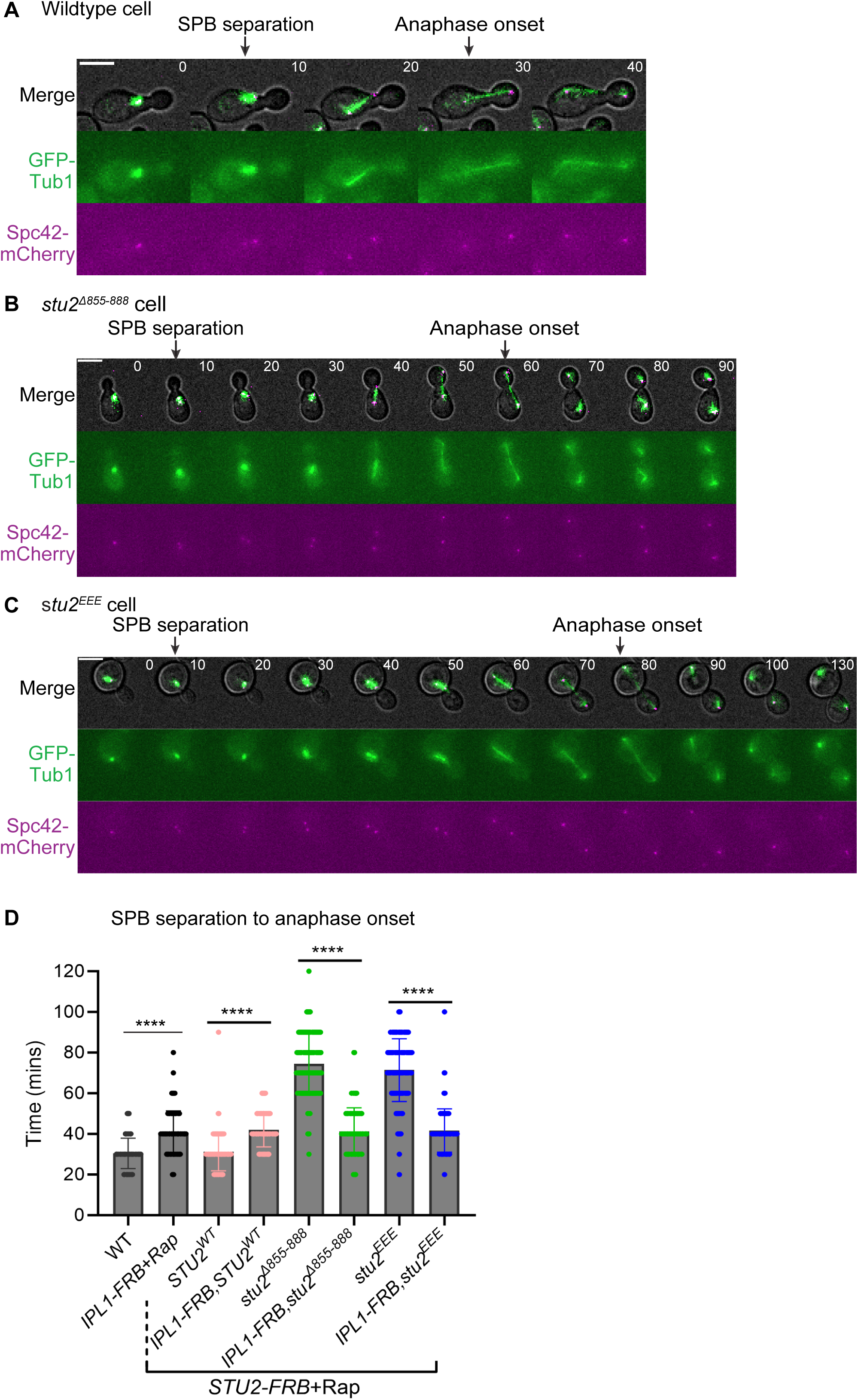
Ipl1 and Stu2 co-depletion accelerates mitotic anaphase onset compared with Stu2 depletion alone. (A-C) Representative mitotic time-lapse images of cells with the indicated genotypes expressing Spc42-mCherry and GFP-Tub1. Scale bars, 5 μm. (D) Graph of the mean time (in minutes) from SPB separation to anaphase onset in cells of the indicated genotypes (****, P < 0.0001; Mann-Whitney test). Rapamycin was added after release from G1 arrest (ɑ-factor wash out). Error bars represent the standard deviation (SD); n ≥100 cells analyzed per genotype.

### Loss of kinetochore-bound Stu2 causes asymmetric chromosome segregation during meiosis

Loss of Ipl1 during both mitosis and meiosis results in extensive chromosome missegregation, with most chromosomes preferentially segregating towards a single spindle pole (Biggins et al., 1999; Tanaka et al., 2002; Cairo et al., 2020; Biggins and Murray, 2001). During budding yeast mitosis, kinetochores remain attached to spindle microtubules throughout most of the cell cycle except for a brief one-two minute period during centromere replication (Kitamura et al., 2007). Following SPB separation, preexisting kinetochore-microtubule attachments are retained by the old SPB, and Ipl1-dependent release of these attachments is required to allow chromosomes to establish bipolar attachments (Dewar et al., 2004; Biggins et al., 1999; Tanaka et al., 2002; Biggins and Murray, 2001). In contrast, during meiosis, outer kinetochore components are shed during prophase I and kinetochore-microtubule attachments are released before they are reestablished upon entry into the meiotic divisions (Meyer et al., 2015; Kim et al., 2013; Miller et al., 2012). During this process, microtubules from the old SPB initially capture more kinetochores than those from the new SPB, likely because the old SPB has nucleated more microtubules at that time (Meyer et al., 2013). Ipl1-dependent release of these initial attachments is therefore required to redistribute chromosomes between the two SPBs and promote biorientation.

We asked whether loss of kinetochore-bound Stu2 results in meiotic chromosome segregation defects similar to those observed following Ipl1 depletion (Cairo et al., 2020, 2023). To address this question, we examined bulk chromosome segregation in cells depleted of Stu2 and/or Ipl1 by monitoring the distribution of fluorescently tagged histone Htb2 (*HTB2-mCherry*). If chromosomes segregate evenly, Htb2-mCherry nuclear masses should be of equal size following meiosis I and II (Figure 4A). In contrast, unequal chromosome segregation is expected to produce Htb2-mCherry masses of different sizes at each meiotic division (Figure 4B).

**Figure 4.**
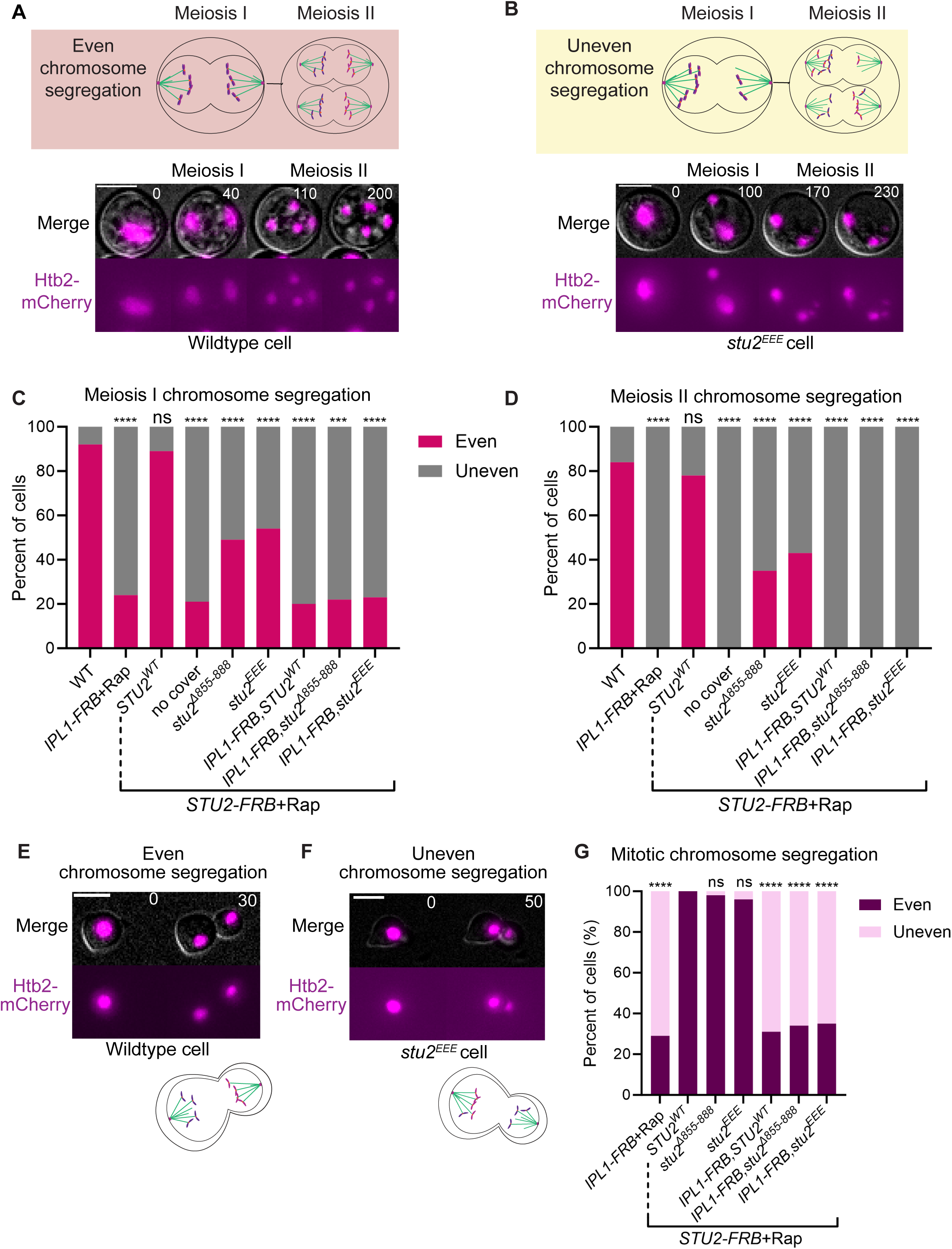
Stu2 kinetochore depletion causes chromosome missegregation in meiosis, with preferential segregation to one SPB. (A-B) Schematic and representative time-lapse images of wildtype cells displaying even chromosome segregation (A) and *stu2^EEE^*cells displaying uneven chromosome segregation (B) in meiosis I and II. Cells express Htb2-mCherry to monitor chromatin. Scale bars, 5 μm. (C and D) Percentage of cells with even and uneven chromosome segregation in meiosis I (C) or meiosis II (D) in the indicated genotypes. Asterisks indicate a statistically significant difference from wildtype (****, P < 0.0001; ns, not significant; two-tailed Fisher’s exact test). n ≥100 cells per genotype. Rap, rapamycin addition. (E and F) Representative mitotic time-lapse images of wildtype mitotic cells displaying even chromosome segregation (E) and Stu2-depleted cells expressing *stu2^EEE^* displaying uneven chromosome segregation (F). Scale bars, 5 μm. (G) Percentage of cells that display even and uneven chromosome segregation in mitosis in the indicated genotypes. Asterisks indicate a statistically significant difference from wildtype cells (**** P < 0.0001; ns, not significant; two-tailed Fisher’s exact test). n ≥100 cells analyzed per genotype. Rap, rapamycin.

Depletion of either Ipl1 or Stu2 significantly increased uneven chromosome segregation during both meiosis I and meiosis II, with many chromosomes segregating toward one spindle pole (Figure 4C-D). The kinetochore-defective *stu2* mutants exhibited a similar phenotype: 50% of *stu2^Δ855-888^* and 45% of *stu2^EEE^* cells displayed uneven chromosome segregation during meiosis I (Figure 4C). Remarkably, all mutants exhibited more severe chromosome segregation defects during meiosis II, which may reflect the combined effects of unequal segregation during both meiosis I and meiosis II (Figure 4D). Co-depletion of Ipl1 and Stu2 in cells expressing *stu2^Δ855-888^* or *stu2^EEE^* further increased the frequency of uneven chromosome segregation, similar to Ipl1 depletion alone (Figure 4C-D). Thus, loss of kinetochore-bound Stu2 causes significant defects in chromosome segregation during meiosis that resembles the phenotype caused by Ipl1 depletion.

These meiotic results prompted us to ask whether loss of kinetochore-bound Stu2 causes a similar chromosome segregation defect during mitosis. As previously reported, Ipl1 depletion caused pronounced uneven chromosome segregation during mitosis, consistent with its critical role in releasing improper kinetochore-microtubule attachments (Dewar et al., 2004; Biggins et al., 1999; Tanaka et al., 2002; Cairo et al., 2020, 2023; Biggins and Murray, 2001) (Figure 4G). In contrast to the severe defects observed during meiosis, loss of kinetochore-bound Stu2 in mitosis did not cause a significant defect (Figure 4G). Co-depletion of Ipl1 and kinetochore-bound Stu2 resulted in more severe segregation defects, resembling those caused by Ipl1 depletion alone. Thus, severe reduction of kinetochore-bound Stu2 has a substantially greater effect on chromosome segregation during meiosis than during mitosis, revealing a heightened requirement for kinetochore-bound Stu2 in meiotic chromosome segregation.

### Cells lacking kinetochore-bound Stu2 segregate chromosomes asymmetrically toward either SPB

Depletion of Ipl1 and loss of kinetochore-bound Stu2 resulted in similar bulk chromosome segregation defects during meiosis (Figure 4G). However, Ipl1 and kinetochore-bound Stu2 differentially regulate kinetochore-microtubule attachments, which suggests that these chromosome segregation defects may arise through distinct mechanisms (Miller et al., 2016, 2019; Zahm et al., 2021). Uneven chromosome segregation following Ipl1 depletion results from a failure to efficiently release initial improper kinetochore-microtubule attachments (Dewar et al., 2004; Biggins et al., 1999; Tanaka et al., 2002; Cairo et al., 2020, 2023; Biggins and Murray, 2001). In contrast, kinetochore-defective *stu2* mutants activate the spindle checkpoint in meiosis, consistent with the presence of unattached kinetochores and indicating that kinetochore-microtubule attachments can be released (Figure 1E-F). We therefore asked whether the asymmetric chromosome segregation in *stu2* mutants is biased, as observed following Ipl1 depletion, or occurs toward either the old or new SPB.

To distinguish between these possibilities, we monitored bulk chromosome segregation (Htb2-GFP) in cells expressing Spc42-RedStar2, a slow-maturing RFP that distinguishes between the old (brighter) and new (dimmer) SPB following spindle pole separation (Meyer et al., 2013; Cairo et al., 2020; Janke et al., 2004)(Figure 5A). In both meiosis I and meiosis II, *stu2* mutant cells that underwent massive chromosome missegregation frequently exhibited the larger chromatin mass at either the old or the new SPBs (Figure 5B-D). This result contrasts with Ipl1-depleted cells, in which the larger chromatin mass is predominantly associated with the old SPB (Meyer et al., 2013; Cairo et al., 2020, 2023)(Figure 5D). Together, these findings indicate that the initial erroneous kinetochore-microtubule attachments are released in *stu2* mutants. However, subsequent reattachment remains highly error prone, allowing either spindle pole to capture a disproportionate number of chromosomes.

**Figure 5.**
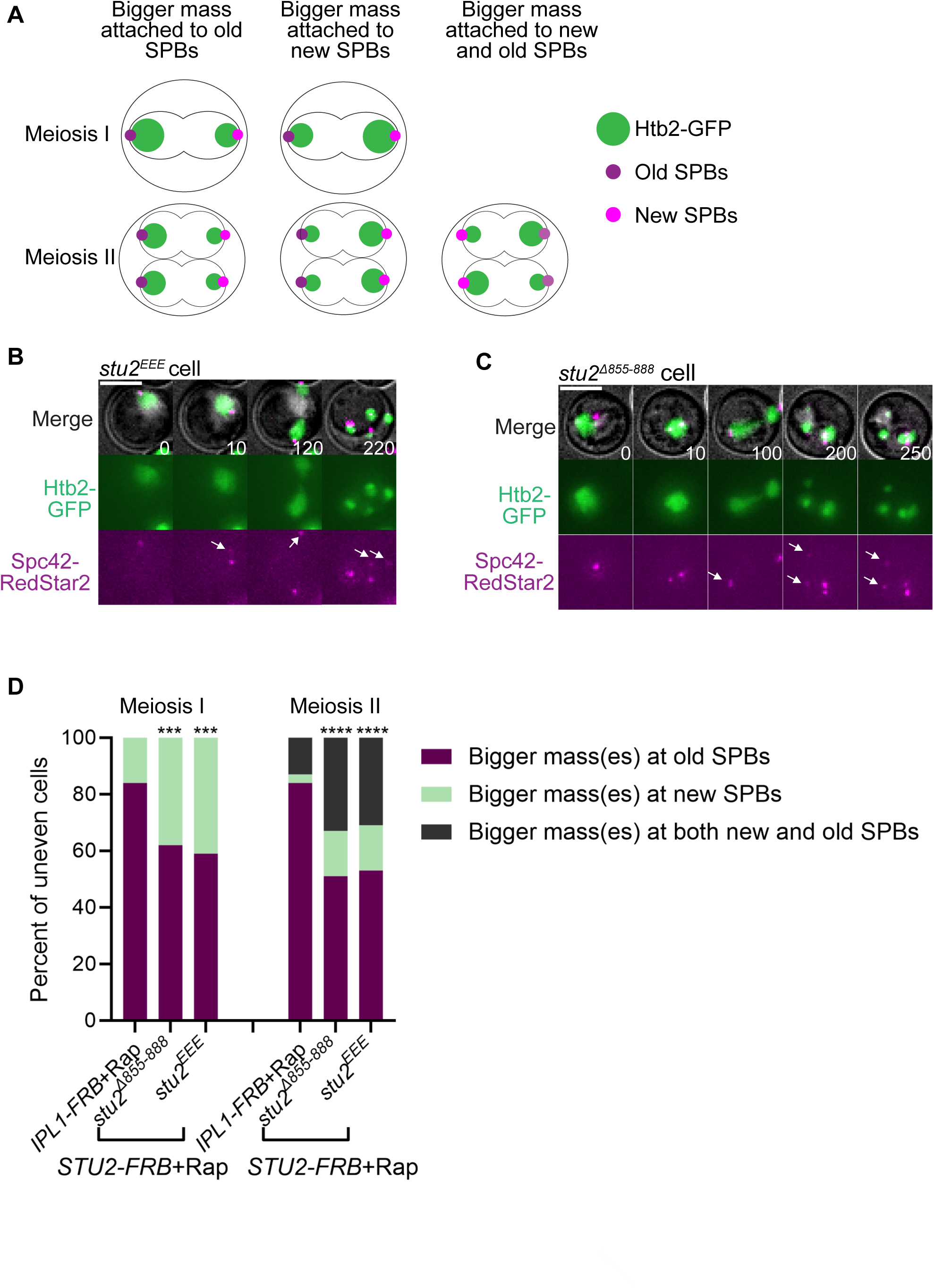
Cells lacking kinetochore-bound Stu2 asymmetrically segregate chromosomes to either the old or new SPB. (A) Schematic showing the attachment of even and uneven chromosome masses at either the dimmer (new) SPBs and brighter (old) SPB. (B-D) Representative time-lapse images of a *stu2^EEE^* (B), or a *stu2^Δ855-888^* cell (C). Cells express Htb2-GFP and Spc42-RedStar2 to monitor DNA masses and SPBs. Scale bars, 5 μm. Arrowheads mark the dimmer (new) SPBs. (D) Percentage of cells in which the biggest DNA masses migrate to either the dimmer (new) SPBs or the brighter (old) SPB in meiosis I and meiosis II. Asterisks indicate a statistically significant difference from Ipl1 depleted cells (***, P-0.0005; ****, P < 0.0001; Chi-square test). n ≥70 cells analyzed per genotype. Rap, rapamycin addition.

### Loss of kinetochore-bound Stu2 causes more frequent chromosome missegregation during meiosis than during mitosis

To further define the role of kinetochore-bound Stu2 during meiotic chromosome segregation, we followed the behavior of an individual chromosome pair. LacO arrays were integrated near the centromere of chromosome IV in cells expressing GFP-LacI, allowing us to track chromosome IV segregation by time-lapse microscopy (Straight et al., 1996) (Figure 6A-B). These cells also express mRuby-Tub1 to monitor spindle microtubules, and images were acquired at five-minute intervals throughout both meiotic divisions (Markus et al., 2015). In the majority of wildtype cells, chromosome IV segregated accurately during both meiosis I and meiosis II (Figure 6B, E-F). In contrast, 34% of *stu2^Δ855-88^*^8^ and 32% of *stu2^EEE^* cells missegregated chromosome IV in meiosis I (Figure 6C-F). The frequency of chromosome IV missegregation was even greater during meiosis II, indicating that additional segregation errors occur during the second meiotic division beyond those inherited from meiosis I (Figure 6F). Furthermore, co-depletion of Ipl1 and kinetochore-bound Stu2 resulted in high frequencies of chromosome IV missegregation during both meiotic divisions, resembling the phenotype caused by an Ipl1 depletion alone (Figure 6E-F).

**Figure 6.**
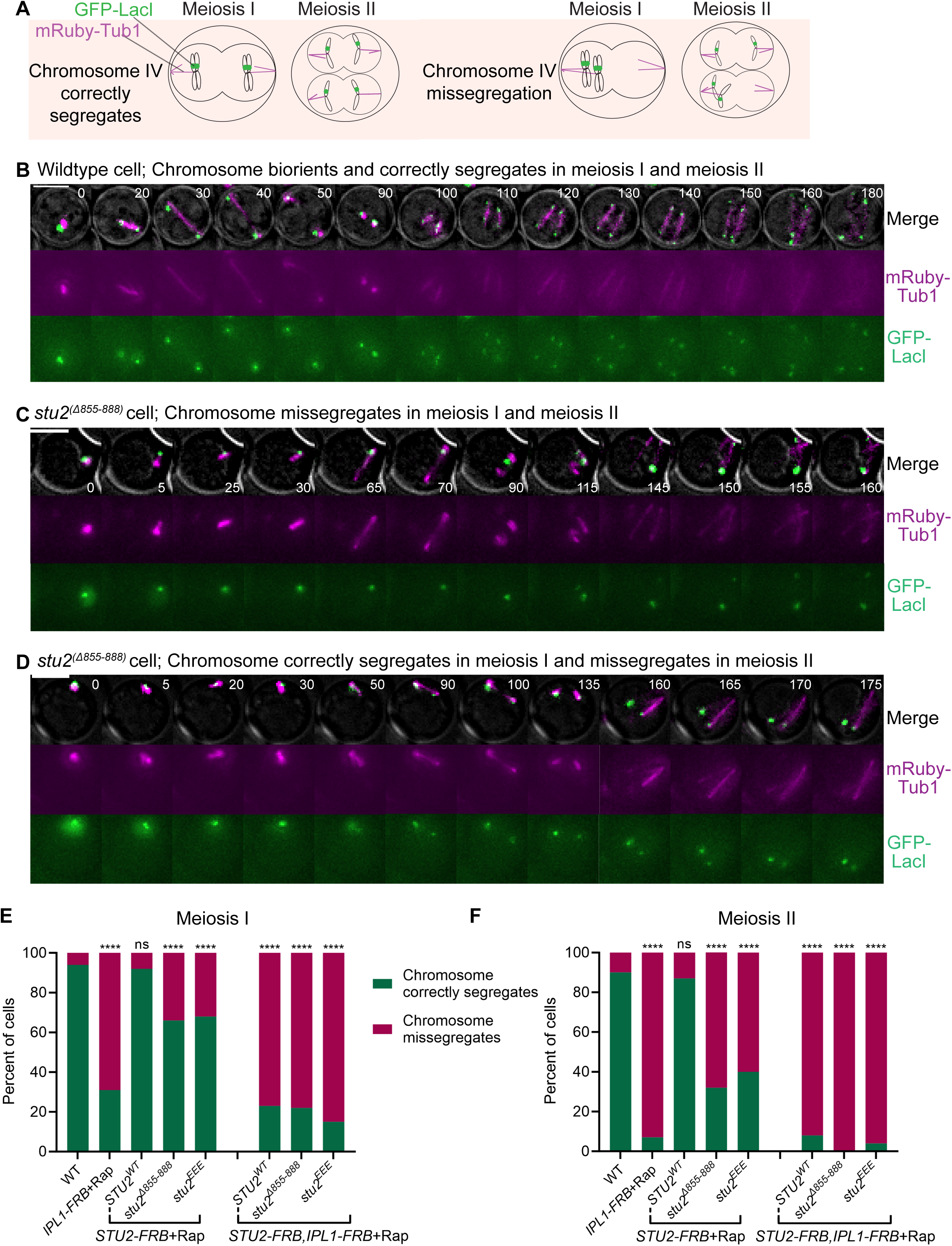
Loss of kinetochore-bound Stu2 increases chromosome missegregation during meiosis. (A) Schematic of chromosome segregation with a LacO/LacI-GFP tagged chromosome IV, with LacO integrated near the centromere. Cells express LacI-GFP and mRuby-Tub1. (B-D) Representative meiotic time-lapse images of a wildtype cell that properly segregates chromosome IV in meiosis I and II (B), a Stu2-depleted cell expressing *stu2^Δ855-888^* showing chromosome IV missegregation in meiosis I and II (C), and a Stu2-depleted cell expressing *stu2^Δ855-888^* showing chromosome IV proper segregation in meiosis I and missegregation in meiosis II (D). Scale bars, 5 μm. (E and F) Graphs of the percent of cells that display proper segregation and missegregation of chromosome IV in meiosis I (E) and meiosis II (F). Asterisks indicate a statistically significant difference from wildtype cells (****, P < 0.0001; ns, not significant; two-tailed Fisher’s exact test). n ≥100 cells analyzed per genotype. Rap, rapamycin.

We next examined whether kinetochore-bound Stu2 is similarly required for accurate chromosome segregation during mitosis. In contrast to meiosis, expression of *stu2^Δ855-888^* and *stu2^EEE^*did not significantly increase chromosome IV missegregation during mitosis (Figure 7A-D). Co-depletion of Ipl1 and kinetochore-bound Stu2 produced a segregation defect similar to Ipl1 depletion alone. Thus, a strong reduction in kinetochore-bound Stu2 causes a minor percent of chromosome IV missegregation in mitosis, but a much higher percent of chromosome missegregation in meiosis. These findings reveal a more sensitive requirement for kinetochore-bound Stu2 in ensuring accurate chromosome segregation during meiosis.

**Figure 7.**
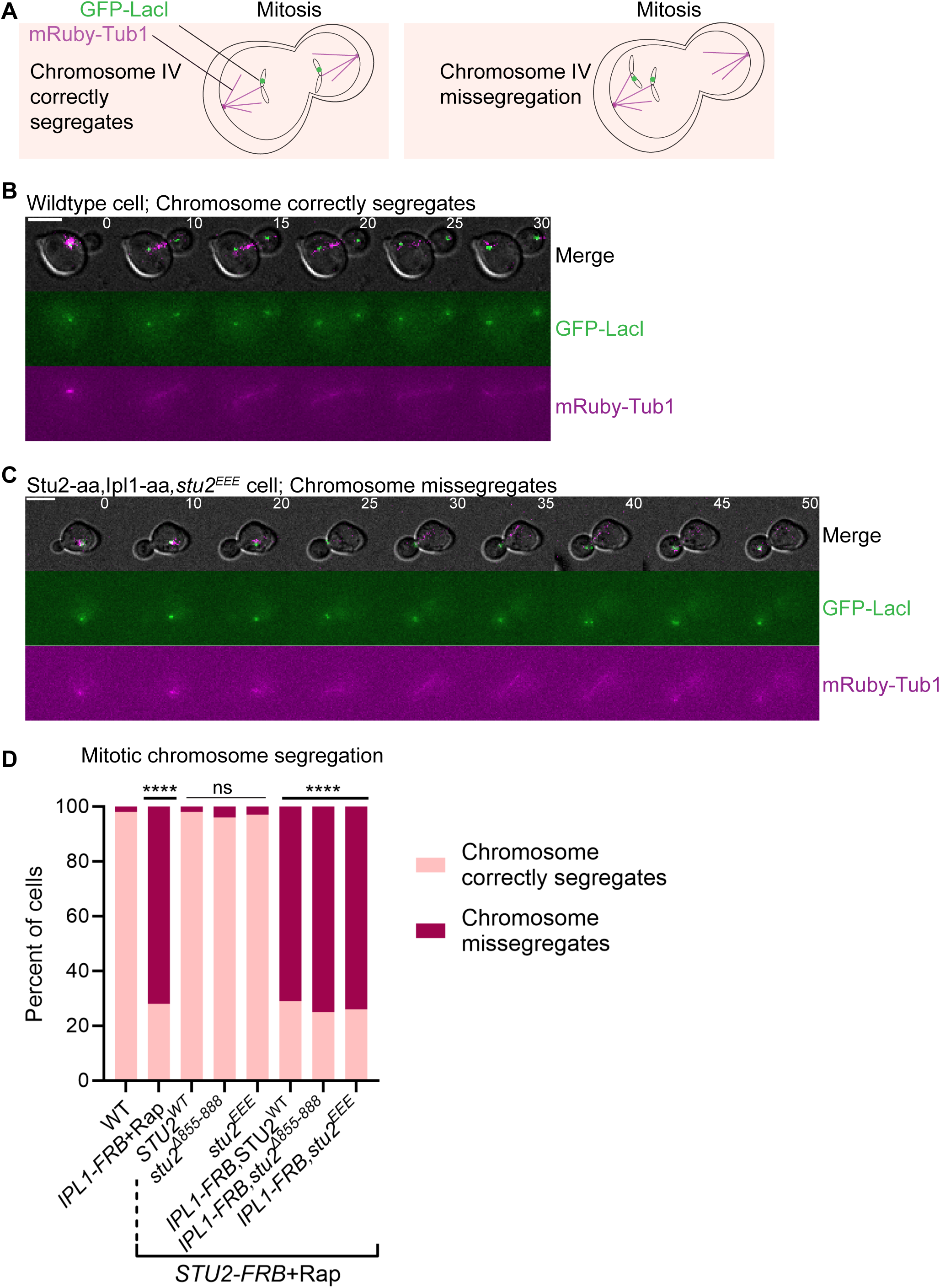
Loss of kinetochore-bound Stu2 only causes modest chromosome missegregation during mitosis unless combined with Ipl1 depletion. (A) Schematic of mitotic chromosome segregation using LacO/LacI-GFP marked chromosome IV. LacO repeats are integrated near the centromere and cells express LacI-GFP and mRuby-Tub1. (B-C) Representative mitotic time-lapse images of a wildtype cell that properly segregates chromosome IV (B), or a Stu2- and Ipl1-depleted cell expressing *stu2^EEE^* that missegregates chromosome IV in mitosis (C). Scale bars, 5 μm. (D) Percent of cells that display proper segregation and missegregation of chromosome IV in mitosis with the indicated genotypes. Asterisks indicate a statistically significant difference from wildtype (****, P < 0.0001; ns, not significant; two-tailed Fisher’s exact test). n ≥100 cells analyzed per genotype. Rap, rapamycin.

### Stu2 kinetochore-defective mutants undergo repeated attempts at error correction, unlike Ipl1-depleted cells

To further define the distinct roles of Stu2 and Ipl1 in chromosome segregation, we monitored LacO/LacI-GFP-labeled chromosome IV dynamics relative to the spindle poles during meiosis by time-lapse microscopy (Figure 8A). Cells were imaged at five-minute intervals, and the positions of GFP-labeled chromosome IV homologs were tracked relative to the two SPBs. In wild-type cells, chromosome IV rapidly achieved biorientation, with the homologs positioned between the two spindle poles, before segregating to opposite poles (Figure 8B). Transient improper orientations, in which both homologs were positioned closer to the same pole were efficiently corrected. In contrast, chromosome IV homologs in Ipl1-depleted cells typically remained associated with a single SPB and rarely traversed the spindle, consistent with previous findings that Ipl1 is required for the release of initial improper kinetochore-microtubule attachments (Meyer et al., 2013; Cairo et al., 2020, 2023) (Figure 8D-F). Strikingly, chromosome IV behaved differently in kinetochore-defective *stu2* mutants. The GFP-labeled chromosomes repeatedly moved between the SPBs and the spindle midzone, consistent with repeated attempts to correct improper attachments (Figure 8C-F). Despite these attempts, chromosome IV frequently missegregated (Figure 6), suggesting that kinetochore-bound Stu2 is required to efficiently establish or stabilize productive attachments. Together, these observations distinguish the roles of Ipl1 and kinetochore-bound Stu2 during meiotic error correction and support a model in which Ipl1 promotes the release of erroneous kinetochore-microtubule attachments, whereas Stu2 promotes the establishment and stabilization of attachments that lead to proper biorientation.

**Figure 8.**
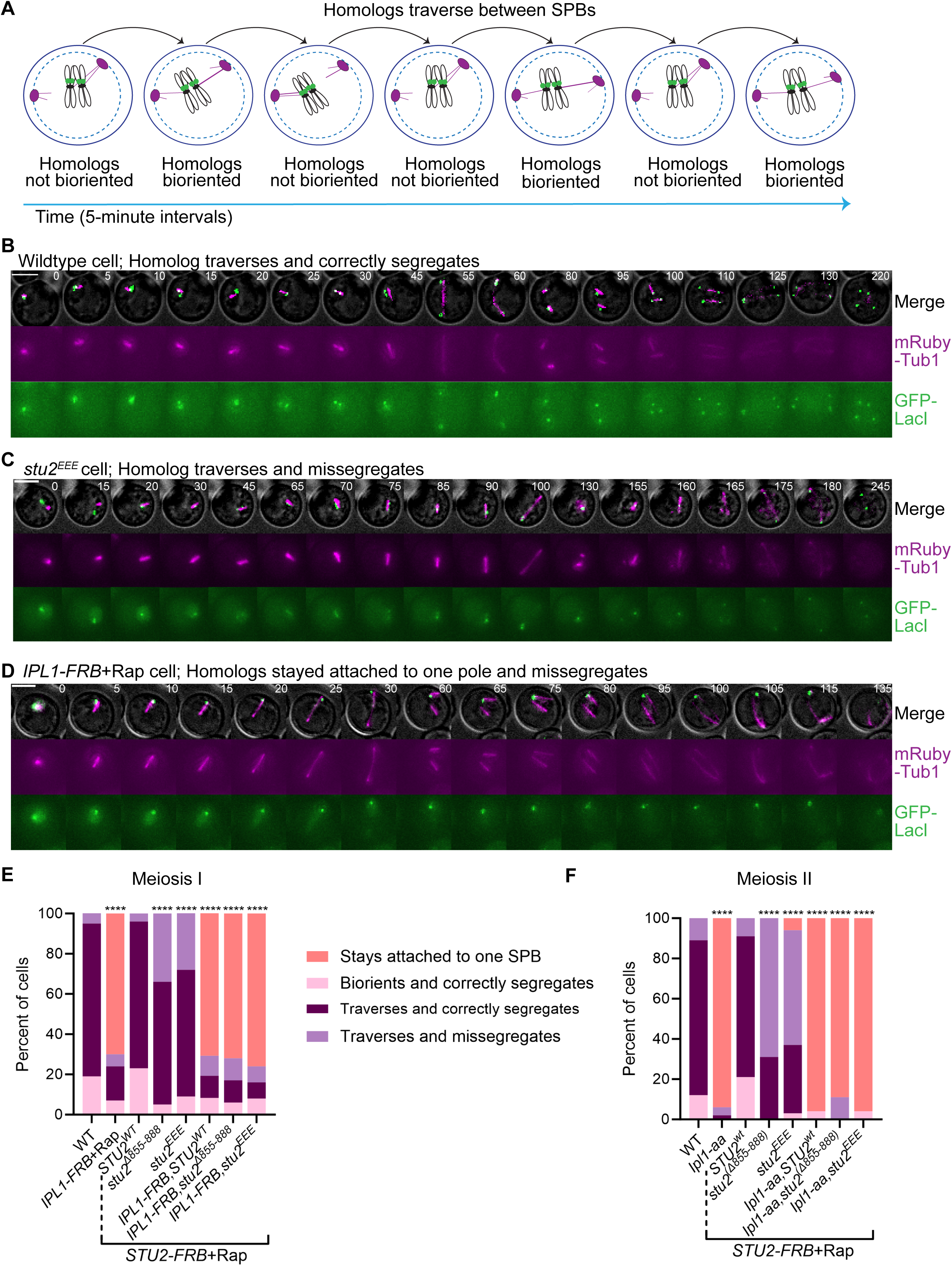
Cells lacking kinetochore-bound Stu2, but not Ipl1-depleted cells, undergo error correction of kinetochore-microtubule attachments. (A) Schematic illustrating the traversal of LacO/LacI-tagged chromosome IV as an indicator of kinetochore-microtubule error correction during chromosome biorientation. LacO repeats are integrated near the centromere of chromosome IV and cells express LacI-GFP and mRuby-Tub1 to monitor spindle microtubules. (B-D) Representative meiotic time-lapse images of a wildtype cell in which chromosome IV undergoes traversal before achieving biorientation and accurate segregation (B), a Stu2-depleted cell expressing *stu2^EEE^* in which chromosome IV undergoes traversal but missegregates (C), or an Ipl1-depleted cell in which chromosome IV fails to undergo traversal and subsequently missegregates (D). Scale bars, 5 μm. (E-F) Quantification of chromosome IV dynamics during error correction, showing the percentage of cells exhibiting chromosome biorientation, traversal, and missegregation during meiosis I (E) and meiosis II (F). Asterisks indicate statistically significant differences from wildtype cells. (**** P < 0.0001; Chi-square test). n ≥100 cells analyzed per genotype. Rap, rapamycin.

## Discussion

Cells have evolved multiple mechanisms to ensure the establishment of tension-bearing kinetochore-microtubule attachments and prevent chromosome missegregation. One such mechanism is the spindle checkpoint, which delays anaphase onset when kinetochores are unattached to spindle microtubules, providing cells additional time to establish the accurate attachments prior to segregation (Bhalla and Lacefield, 2026; Pleuger and Westermann, 2026). In parallel, multiple error correction pathways destabilize improper attachments that fail to generate tension while promoting the stabilization of tension-bearing attachments (Li et al., 2024). These pathways ensure that kinetochores can undergo multiple rounds of microtubule release and reattachment until tension-bearing attachments are established.

Here, we compared the Stu2^chTOG^ and Ipl1^Aurora^ ^B^ error correction pathways in both meiosis and mitosis in budding yeast. Stu2 has a well-established role as a microtubule polymerase, but the essential role is in error correction of kinetochore-microtubule attachments (Li et al., 2024; Abouelghar et al., 2025). The Stu2 C-terminal segment was previously shown to bind the four-way junction of the Ndc80 complex, interacting with Ndc80 and Spc24 (Zahm et al., 2021). We analyzed two different mutants each with substantially diminished binding of Stu2 to the kinetochore receptors within the Ndc80 complex, *stu2^Δ855-888^*and *stu2^EEE^* (Zahm et al., 2021). Both *stu2^Δ855-888^*and *stu2^EEE^* abrogate Ndc80 complex interactions and have similar phenotypes in our assays.

Our analysis revealed important differences in the requirement for kinetochore-bound Stu2 between meiosis and mitosis. Notably, a substantial loss of kinetochore-bound Stu2 results in more severe phenotypes during meiosis than in mitosis. First, *stu2^Δ855-888^* and *stu2^EEE^* cells exhibited a longer spindle checkpoint delay in meiosis than in mitosis (Figures 1E-F and 3D). This difference suggests that *stu2* mutants may either generate more erroneous kinetochore-microtubule attachments during meiosis or establish new attachments less efficiently following their release. Second, we found that a large fraction of *stu2^Δ855-888^* and *stu2^EEE^* cells exhibited asymmetric chromosome segregation in both meiosis I and meiosis II, with an excess of chromosomes segregating toward one SPB (Figure 4). This phenotype was observed at a substantially lower frequency in mitotic *stu2^Δ855-888^* and *stu2^EEE^* cells. In meiosis, the excess of chromosomes segregated toward either the old or new SPB, indicating that the asymmetry was not due to a failure to release initial attachments (Figure 5). Third, analysis of an individual chromosome revealed a much higher frequency of chromosome missegregation in meiosis than in mitosis (Figure 6 and 7). Together, these findings indicate that full kinetochore-bound Stu2 is particularly important for ensuring accurate chromosome segregation during meiosis and suggest a greater requirement for Stu2-mediated error correction in meiosis than in in mitosis.

Why meiotic error correction depends on a much higher level of kinetochore-bound Stu2 than mitotic error correction remains an interesting question. Interestingly, Mps1 mutants also exhibit more severe chromosome missegregation in meiosis than in mitosis. We propose that differences in kinetochore architecture, cohesin linkages, and cell-cycle duration may make meiotic kinetochore-microtubule attachment particularly error-prone, increasing the requirement for multiple error correction pathways. In addition, alternative interactions may support Stu2 function at kinetochores when its primary kinetochore-binding interface is disrupted. Previous work has shown that efficient Stu2 kinetochore association also depends on its TOG1 domain and homodimerization, while interactions with other kinetochore proteins, such as Cnn1, may provide additional modes of kinetochore association (Humphrey et al., 2018; Miller et al., 2019; Abouelghar et al., 2025; Maitra et al., 2026). These weaker or alternative interactions may provide sufficient Stu2 activity to support error correction during mitosis but may be inadequate to meet the greater demands of chromosome biorientation during meiosis.

Our comparison of *stu2^Δ855-888^* and *stu2^EEE^* cells to Ipl1-depleted cells revealed several important differences between the functions of Stu2 and Ipl1 during meiosis. Loss of kinetochore-bound Stu2 resulted in a strong spindle checkpoint-dependent delay, whereas Ipl1-depleted cells did not activate the spindle checkpoint, consistent with their inability to efficiently release initial improper kinetochore-microtubule attachments (Figure 1). Both Stu2 kinetochore-defective and Ipl1-depleted cells also exhibited asymmetric chromosome segregation, with an excess of chromosomes segregating toward one spindle pole (Figure 4). Remarkably, however, *stu2^Δ855-888^* or *stu2^EEE^* cells segregated excess chromosomes toward either the old or the new SPB, whereas Ipl1-depleted cells preferentially segregated the majority of chromosomes toward the old SPB (Meyer et al., 2013; Cairo et al., 2020, 2023) (Figure 5). These results suggest that although initial attachments are likely released, kinetochore-defective Stu2 cells are unable to ensure equal chromosome segregation to both poles. Analysis of a single tagged chromosome further revealed that cells with kinetochore-depleted Stu2 undergo attempts at error correction, whereas most Ipl1-depleted cells fail to release their initial attachments (Figure 6 and 8). Finally, co-depletion of Ipl1 and kinetochore-bound Stu2 produced phenotypes resembling Ipl1 depletion alone (Figure 2-4, 6-8). Together, these findings support distinct roles for Ipl1 and Stu2 in meiotic error correction where Ipl1 promotes the release of initial erroneous attachments and Stu2 promotes the subsequent stabilization of correct attachments and possibly further release of unstable attachments. Importantly, when preexisting kinetochore-microtubule attachments are disrupted with microtubule-depolymerizing drugs, chromosomes segregate randomly in Ipl1-depleted cells, suggesting that Ipl1 has additional roles in establishing biorientation beyond releasing initial attachments (Monje-Casas et al., 2007).

Our analyses of Stu2 kinetochore-deficient mutants in meiosis, together with previous studies of Ipl1 and Mps1, provide a framework for distinguishing the contributions of these three error correction pathways. Previous work demonstrated that Ipl1 and Mps1 function sequentially during meiotic error correction: following Ipl1-dependent release of improper attachments, Mps1 promotes the formation of new force-generating kinetochore-microtubule attachments (Meyer et al., 2013). Although both Mps1 and Stu2 promote the stabilization of tension-bearing attachments, they likely do so through distinct mechanisms. Mps1 promotes the formation and stabilization of force-generating end-on kinetochore-microtubule attachments, whereas *in vitro* studies have shown that Stu2 preferentially stabilizes attachments to assembling microtubule tips under tension while destabilizing low tension attachments at disassembling tips (Meyer et al., 2018; Miller et al., 2016; Meyer et al., 2013, 2021, 2015). Altogether, these findings support a model in which Ipl1, Mps1, and Stu2 provide complementary activity during meiotic error correction: Ipl1 promotes the release of improper attachments, Mps1 facilitates the establishment of new force-generating attachments, and kinetochore-bound Stu2 helps select and stabilize productive attachments while allowing additional rounds of error correction when proper biorientation is not achieved.

## Materials and Methods

### Budding yeast strains and manipulations

All *S. cerevisiae* strains used in this study are W303 derivatives and are listed in Supplemental Table S1. All yeast strains contain the following genetic background: *ade2-1 his3-11, 15 leu2-3, trp1-1 ura3-1*, and *can1-100*. Gene deletions and gene tagging were performed using standard PCR-based amplification of the inserts followed by lithium acetate transformation (Janke et al., 2004). Genetic manipulations were checked by PCR and microscopy for fluorescently tagged genes. Plasmids containing the *stu2* alleles (*STU2^WT^*, *stu2^Δ855-888^*, *stu2^EEE^*) were gifts from Matthew Miller and were integrated at the *LEU2* locus. Strains with the anchor-away background (*tor1-1, fpr1Δ, RPL13-2XFKBP12*) were previously described, with added *STU2* and/or *IPL1* C-terminally tagged with *FRB* at the endogenous locus and *RPL13A* with *2XFKBP12* (Haruki et al., 2008).

### Budding yeast growth conditions

For meiotic experiments, strains were grown in 2X synthetic complete medium (2XSC; 1.34% bacto–yeast nitrogen base without amino acids, 0.4% dropout mix with all amino acids, and 4% glucose) for 18-24 hours at 30°C, transferred with a 1:25 dilution to 1X synthetic complete acetate medium (1XSCA; 0.67% bacto–yeast nitrogen base without amino acids, 0.2% dropout mix with all amino acids, 2% potassium acetate) for 12–16 hour at 30°C, washed three times with water, and then incubated in 1% potassium acetate at 25°C for 7-8 hours to allow cells to reach prophase I. In all experiments with anchor away nuclear depletion, rapamycin (1 μg/ml) was added at the same time cells were transferred to potassium acetate, except for Ipl1 depletion, which was added 7 hours after cells were transferred to potassium acetate. For synchronized mitotic experiments, cells were grown in 2XSC medium overnight at 30°C. These cells were then diluted (1:10) in 2XSC medium and grown for 1 hour at 30°C. 25 μM α-factor was added, and cells were grown for 2-3 hours at 30°C. After checking G1 arrest (unbudded cells) using microscopy, cells were washed 5-6 with 2XSC medium and resuspended in 2XSC medium. In all mitotic experiments with anchor away nuclear depletion, rapamycin (1μg/ml) was added after α-factor wash out. Cells were then grown for 30 minutes at 30°C to allow nuclear depletion, and then time-lapse microscopy was initiated. All cultures were kept on a roller drum at the indicated temperature. All liquid media were supplemented with 1% tryptophan and 0.5% adenine. All reagents used in this study are listed in Supplemental Table S2.

### Microscope image acquisition and time-lapse microscopy

All time-lapse imaging was performed in a chamber mounted on a coverslip coated with Concanavalin A (conA). In all meiosis movies, after 7–8 hours of transferring cells to potassium acetate, 200µl of the cells were concentrated, and 6µL was loaded onto the conA-coated coverslip in a chamber. A 5% agar pad in KAc (or 2XSC for mitotic cells) was made by cutting off the tip of an Eppendorf tube and pipetting ∼100 μL of melted 5% agar into the Eppendorf tube placed on a glass slide/coverslip. The solidified agar pad was then placed on top of the cells for 12 minutes, to allow the cells to adhere to the ConA and to create a monolayer of cells. The rest of the preconditioned 1% KAc (or 2XSC for mitotic cells) culture (∼ 2 mL) was spun down, and the supernatant was added dropwise to the chamber to float and remove the agar pad. The chamber was immediately placed on the microscope for imaging.

Images were acquired on a Nikon Ti2 microscope equipped with a Photometrics camera and 60X inverted-objective microscope equipped with a 60X oil objective. During time-lapse imaging, Z-stacks of five steps of 1.2μm each were acquired at 10-minute intervals (or 5-minute intervals for LacI-LacO strains) with exposure times of 20–50ms for brightfield and 30-50ms for GFP, mCherry or mRuby with neutral-density filters transmitting 2–3% of light intensity. All meiosis movies were set up for 12 hours, while mitosis movies were 4-5 hours long. For each experiment, 100 cells were counted from two or more movies for each genotype, unless otherwise noted.

### Image processing

All microscopy images were collected on the Nikon Ti2 microscope with Z-stacks of five steps of 1.2μm each at 10-minute intervals (or 5-minute intervals for LacI-LacO strains). NIS elements software was used to combine the z-stacks into a single maximum intensity projection for analysis. The images were processed using ImageJ/Fiji software (National Institutes of Health) to create final time-lapse images with adjustment of brightness and contrast. For measuring DNA mass area in mitosis, a circle was drawn around the Htb2-mCherry signal, area was measured, and the two DNA masses were subtracted from each other to obtain a mass size difference. For each individual cell, any mass size difference outside the defined range of the wild type was classified as “uneven.”

### Spotting assay

For spot assays, 10-fold serially dilutions of saturated overnight cultures in YPD were spotted onto YPD plates (1% yeast extract, 2% peptone, 2% glucose) or YPD supplemented with 1μg/ml rapamycin. Plates were incubated at 23°C for 36–40 hours before imaging.

### Statistical analysis

Statistical analysis was performed using Prism (GraphPad Software). Statistical analysis for meiosis and mitosis timings were analyzed using an unpaired, nonparametric Mann-Whitney test with computation of two-tailed exact P values. Statistical analysis of DNA masses (Htb2-mCherry strains) were analyzed using the two-sided Fisher’s exact test. The number of data points (n) is indicated in the figure legends. Statistically significant differences (P value <0.05) are indicated with an asterisk.

## Acknowledgements

We thank Matthew Miller for strains and both Matthew Miller and Ahmed Abouelghar for insightful discussions. We thank the Lacefield Lab and Mackenzie Flynn for critical reading of the manuscript. This work is supported through NIGMS R01GM105755 to SL and through our core facilities, the bioMT P20-GM113132 and the Dartmouth Cancer Center Shared Resource P30-CA023108.

## Figure Legends

**Figure S1. Stu2 kinetochore depletion reduces cell viability.** (A) Ten-fold serial dilutions of the indicated strains were spotted onto YPD plates or YPD plates with 1μg/ml of rapamycin. The plates were incubated at 23°C for 36–40 hours before imaging.

**Table S1. Budding yeast strains used in this study.** Strains are derivatives of W303.

**Table S2. Reagents used in this study.**

**Table S3. Plasmids and Primers used in this study.**

## Notes

### Competing Interest Statement

The authors have declared no competing interest.

## References

1. Abouelghar, A.A., J.S. Carrier, J.R. Torvi, E. Jenson, C. Jones, B. Gangadharan, E.G. Prinslow, L.M. Rice, B. Lagesse, G. Barnes, and M.P. Miller. 2025. Stu2 performs an essential kinetochore function independent of its microtubule polymerase activity. J Cell Biol. 224:e202209025. doi:10.1083/jcb.202209025.

2. Benzi, G., A. Camasses, Y. Atsunori, Y. Katou, K. Shirahige, and S. Piatti. 2020. A common molecular mechanism underlies the role of Mps1 in chromosome biorientation and the spindle assembly checkpoint. EMBO Rep. 21:e50257. doi:10.15252/embr.202050257.

3. Bhalla, N., and S. Lacefield. 2026. Tuning Meiotic Checkpoints: Evolutionary Costs and Mechanistic Trade-Offs. Annu Rev Genet. doi:10.1146/annurev-genet-031326-012917.

4. Biggins, S., and A.W. Murray. 2001. The budding yeast protein kinase Ipl1/Aurora allows the absence of tension to activate the spindle checkpoint. Genes Dev. 15:3118–3129. doi:10.1101/gad.934801.

5. Biggins, S., F.F. Severin, N. Bhalla, I. Sassoon, A.A. Hyman, and A.W. Murray. 1999. The conserved protein kinase Ipl1 regulates microtubule binding to kinetochores in budding yeast. Genes Dev. 13:532–544.

6. Brouhard, G.J., J.H. Stear, T.L. Noetzel, J. Al-Bassam, K. Kinoshita, S.C. Harrison, J. Howard, and A.A. Hyman. 2008. XMAP215 is a processive microtubule polymerase. Cell. 132:79–88. doi:10.1016/j.cell.2007.11.043.

7. Cairo, G., C. Greiwe, G.I. Jung, C. Blengini, K. Schindler, and S. Lacefield. 2023. Distinct Aurora B pools at the inner centromere and kinetochore have different contributions to meiotic and mitotic chromosome segregation. Mol Biol Cell. 34:ar43. doi:10.1091/mbc.E23-01-0014.

8. Cairo, G., and S. Lacefield. 2020. Establishing correct kinetochore-microtubule attachments in mitosis and meiosis. Essays Biochem. 64:277–287. doi:10.1042/EBC20190072.

9. Cairo, G., A.M. MacKenzie, and S. Lacefield. 2020. Differential requirement for Bub1 and Bub3 in regulation of meiotic versus mitotic chromosome segregation. J Cell Biol. 219:e201909136. doi:10.1083/jcb.201909136.

10. Chan, C.S., and D. Botstein. 1993. Isolation and characterization of chromosome-gain and increase-in-ploidy mutants in yeast. Genetics. 135:677–691. doi:10.1093/genetics/135.3.677.

11. Cheeseman, I.M., J.S. Chappie, E.M. Wilson-Kubalek, and A. Desai. 2006. The Conserved KMN Network Constitutes the Core Microtubule-Binding Site of the Kinetochore. Cell. 127:983–997. doi:10.1016/j.cell.2006.09.039.

12. Cullen, C.F., P. Deák, D.M. Glover, and H. Ohkura. 1999. mini spindles: A Gene Encoding a Conserved Microtubule-Associated Protein Required for the Integrity of the Mitotic Spindle in Drosophila. J Cell Biol. 146:1005–1018. doi:10.1083/jcb.146.5.1005.

13. DeLuca, J.G., W.E. Gall, C. Ciferri, D. Cimini, A. Musacchio, and E.D. Salmon. 2006. Kinetochore Microtubule Dynamics and Attachment Stability Are Regulated by Hec1. Cell. 127:969–982. doi:10.1016/j.cell.2006.09.047.

14. Dewar, H., K. Tanaka, K. Nasmyth, and T.U. Tanaka. 2004. Tension between two kinetochores suffices for their bi-orientation on the mitotic spindle. Nature. 428:93–97. doi:10.1038/nature02328.

15. Gard, D.L., and M.W. Kirschner. 1987. A microtubule-associated protein from Xenopus eggs that specifically promotes assembly at the plus-end. J Cell Biol. 105:2203–2215. doi:10.1083/jcb.105.5.2203.

16. Haruki, H., J. Nishikawa, and U.K. Laemmli. 2008. The anchor-away technique: rapid, conditional establishment of yeast mutant phenotypes. Mol Cell. 31:925–932. doi:10.1016/j.molcel.2008.07.020.

17. Herman, J.A., M.P. Miller, and S. Biggins. 2020. chTOG is a conserved mitotic error correction factor. eLife. 9:e61773. doi:10.7554/eLife.61773.

18. Hewitt, L., A. Tighe, S. Santaguida, A.M. White, C.D. Jones, A. Musacchio, S. Green, and S.S. Taylor. 2010. Sustained Mps1 activity is required in mitosis to recruit O-Mad2 to the Mad1-C-Mad2 core complex. J Cell Biol. 190:25–34. doi:10.1083/jcb.201002133.

19. Humphrey, L., I. Felzer-Kim, and A.P. Joglekar. 2018. Stu2 acts as a microtubule destabilizer in metaphase budding yeast spindles. Mol Biol Cell. 29:247–255. doi:10.1091/mbc.E17-08-0494.

20. Janke, C., M.M. Magiera, N. Rathfelder, C. Taxis, S. Reber, H. Maekawa, A. Moreno-Borchart, G. Doenges, E. Schwob, E. Schiebel, and M. Knop. 2004. A versatile toolbox for PCR-based tagging of yeast genes: new fluorescent proteins, more markers and promoter substitution cassettes. Yeast. 21:947–962. doi:10.1002/yea.1142.

21. Jelluma, N., A.B. Brenkman, N.J.F. van den Broek, C.W.A. Cruijsen, M.H.J. van Osch, S.M.A. Lens, R.H. Medema, and G.J.P.L. Kops. 2008. Mps1 phosphorylates Borealin to control Aurora B activity and chromosome alignment. Cell. 132:233–246. doi:10.1016/j.cell.2007.11.046.

22. Jones, M.H., B.J. Huneycutt, C.G. Pearson, C. Zhang, G. Morgan, K. Shokat, K. Bloom, and M. Winey. 2005. Chemical genetics reveals a role for Mps1 kinase in kinetochore attachment during mitosis. Curr Biol. 15:160–165. doi:10.1016/j.cub.2005.01.010.

23. Kakui, Y., M. Sato, N. Okada, T. Toda, and M. Yamamoto. 2013. Microtubules and Alp7-Alp14 (TACC-TOG) reposition chromosomes before meiotic segregation. Nat Cell Biol. 15:786–796. doi:10.1038/ncb2782.

24. Kim, S., R. Meyer, H. Chuong, and D.S. Dawson. 2013. Dual mechanisms prevent premature chromosome segregation during meiosis. Genes Dev. 27:2139–2146. doi:10.1101/gad.227454.113.

25. Kitamura, E., K. Tanaka, Y. Kitamura, and T.U. Tanaka. 2007. Kinetochore microtubule interaction during S phase in Saccharomyces cerevisiae. Genes Dev. 21:3319–3330. doi:10.1101/gad.449407.

26. Kitamura, E., K. Tanaka, S. Komoto, Y. Kitamura, C. Antony, and T.U. Tanaka. 2010. Kinetochores generate microtubules with distal plus ends: their roles and limited lifetime in mitosis. Dev Cell. 18:248–259. doi:10.1016/j.devcel.2009.12.018.

27. Kosco, K.A., C.G. Pearson, P.S. Maddox, P.J. Wang, I.R. Adams, E.D. Salmon, K. Bloom, and T.C. Huffaker. 2001. Control of microtubule dynamics by Stu2p is essential for spindle orientation and metaphase chromosome alignment in yeast. Mol Biol Cell. 12:2870–2880. doi:10.1091/mbc.12.9.2870.

28. Leça, N., F. Barbosa, S. Rodriguez-Calado, A. Esposito Verza, M. Moura, P.D. Pedroso, I. Pinto, E. Artes, T. Bange, C.E. Sunkel, M. Barisic, T.J. Maresca, and C. Conde. 2025. Proximity-based activation of AURORA A by MPS1 potentiates error correction. Curr Biol. 35:1935–1947.e8. doi:10.1016/j.cub.2025.03.018.

29. Li, S., T. Kasciukovic, and T.U. Tanaka. 2024. Kinetochore-microtubule error correction for biorientation: lessons from yeast. Biochem Soc Trans. 52:29–39. doi:10.1042/BST20221261.

30. Maciejowski, J., H. Drechsler, K. Grundner-Culemann, E.R. Ballister, J.-A. Rodriguez-Rodriguez, V. Rodriguez-Bravo, M.J.K. Jones, E. Foley, M.A. Lampson, H. Daub, A.D. McAinsh, and P.V. Jallepalli. 2017. Mps1 Regulates Kinetochore-Microtubule Attachment Stability via the Ska Complex to Ensure Error-Free Chromosome Segregation. Dev Cell. 41:143–156.e6. doi:10.1016/j.devcel.2017.03.025.

31. Maciejowski, J., K.A. George, M.-E. Terret, C. Zhang, K.M. Shokat, and P.V. Jallepalli. 2010. Mps1 directs the assembly of Cdc20 inhibitory complexes during interphase and mitosis to control M phase timing and spindle checkpoint signaling. J Cell Biol. 190:89–100. doi:10.1083/jcb.201001050.

32. Maitra, N., D.T. Edwards, C. Hu, C.L. Asbury, and S. Biggins. 2026. Kinetochore-microtubule attachments are strengthened by Cnn1 stabilization of Stu2. 2026.07.13.738289. doi:10.64898/2026.07.13.738289.

33. Markus, S.M., S. Omer, K. Baranowski, and W.-L. Lee. 2015. Improved Plasmids for Fluorescent Protein Tagging of Microtubules in Saccharomyces cerevisiae. Traffic. 16:773–786. doi:10.1111/tra.12276.

34. Maure, J.-F., E. Kitamura, and T.U. Tanaka. 2007. Mps1 kinase promotes sister-kinetochore bi-orientation by a tension-dependent mechanism. Curr Biol. 17:2175–2182. doi:10.1016/j.cub.2007.11.032.

35. McVey, S.L., J.K. Cosby, and N.J. Nannas. 2021. Aurora B Tension Sensing Mechanisms in the Kinetochore Ensure Accurate Chromosome Segregation. Int J Mol Sci. 22:8818. doi:10.3390/ijms22168818.

36. Meyer, R.E., J. Brown, L. Beck, and D.S. Dawson. 2018. Mps1 promotes chromosome meiotic chromosome biorientation through Dam1. Mol Biol Cell. 29:479–489. doi:10.1091/mbc.E17-08-0503.

37. Meyer, R.E., H.H. Chuong, M. Hild, C.L. Hansen, M. Kinter, and D.S. Dawson. 2015. Ipl1/Aurora-B is necessary for kinetochore restructuring in meiosis I in Saccharomyces cerevisiae. Mol Biol Cell. 26:2986–3000. doi:10.1091/mbc.E15-01-0032.

38. Meyer, R.E., S. Kim, D. Obeso, P.D. Straight, M. Winey, and D.S. Dawson. 2013. Mps1 and Ipl1/Aurora B act sequentially to correctly orient chromosomes on the meiotic spindle of budding yeast. Science. 339:1071–1074. doi:10.1126/science.1232518.

39. Meyer, R.E., A.R. Tipton, R. LaVictoire, G.J. Gorbsky, and D.S. Dawson. 2021. Mps1 promotes poleward chromosome movements in meiotic prometaphase. Mol Biol Cell. 32:1020–1032. doi:10.1091/mbc.E20-08-0525-T.

40. Miller, M.P., C.L. Asbury, and S. Biggins. 2016. A TOG Protein Confers Tension Sensitivity to Kinetochore-Microtubule Attachments. Cell. 165:1428–1439. doi:10.1016/j.cell.2016.04.030.

41. Miller, M.P., R.K. Evans, A. Zelter, E.A. Geyer, M.J. MacCoss, L.M. Rice, T.N. Davis, C.L. Asbury, and S. Biggins. 2019. Kinetochore-associated Stu2 promotes chromosome biorientation in vivo. PLoS Genet. 15:e1008423. doi:10.1371/journal.pgen.1008423.

42. Miller, M.P., E. Unal, G.A. Brar, and A. Amon. 2012. Meiosis I chromosome segregation is established through regulation of microtubule-kinetochore interactions. Elife. 1:e00117. doi:10.7554/eLife.00117.

43. Monje-Casas, F., V.R. Prabhu, B.H. Lee, M. Boselli, and A. Amon. 2007. Kinetochore orientation during meiosis is controlled by Aurora B and the monopolin complex. Cell. 128:477–490. doi:10.1016/j.cell.2006.12.040.

44. Pleuger, R., and S. Westermann. 2026. Mitotic error correction and the spindle assembly checkpoint: a tension-filled relationship. Cell Cycle. 25:1–19. doi:10.1080/15384101.2026.2653527.

45. Podolski, M., M. Mahamdeh, and J. Howard. 2014. Stu2, the Budding Yeast XMAP215/Dis1 Homolog, Promotes Assembly of Yeast Microtubules by Increasing Growth Rate and Decreasing Catastrophe Frequency *. Journal of Biological Chemistry. 289:28087–28093. doi:10.1074/jbc.M114.584300.

46. Santaguida, S., A. Tighe, A.M. D’Alise, S.S. Taylor, and A. Musacchio. 2010. Dissecting the role of MPS1 in chromosome biorientation and the spindle checkpoint through the small molecule inhibitor reversine. J Cell Biol. 190:73–87. doi:10.1083/jcb.201001036.

47. Sarangapani, K.K., L.B. Koch, C.R. Nelson, C.L. Asbury, and S. Biggins. 2021. Kinetochore-bound Mps1 regulates kinetochore-microtubule attachments via Ndc80 phosphorylation. J Cell Biol. 220:e202106130. doi:10.1083/jcb.202106130.

48. Shirasu-Hiza, M., P. Coughlin, and T. Mitchison. 2003. Identification of XMAP215 as a microtubule-destabilizing factor in Xenopus egg extract by biochemical purification. J Cell Biol. 161:349–358. doi:10.1083/jcb.200211095.

49. Straight, A.F., A.S. Belmont, C.C. Robinett, and A.W. Murray. 1996. GFP tagging of budding yeast chromosomes reveals that protein-protein interactions can mediate sister chromatid cohesion. Curr Biol. 6:1599–1608. doi:10.1016/s0960-9822(02)70783-5.

50. Tanaka, T.U., N. Rachidi, C. Janke, G. Pereira, M. Galova, E. Schiebel, M.J.R. Stark, and K. Nasmyth. 2002. Evidence that the Ipl1-Sli15 (Aurora kinase-INCENP) complex promotes chromosome bi-orientation by altering kinetochore-spindle pole connections. Cell. 108:317–329. doi:10.1016/s0092-8674(02)00633-5.

51. Usui, T., H. Maekawa, G. Pereira, and E. Schiebel. 2003. The XMAP215 homologue Stu2 at yeast spindle pole bodies regulates microtubule dynamics and anchorage. EMBO J. 22:4779–4793. doi:10.1093/emboj/cdg459.

52. Vukušić, K., and I.M. Tolić. 2022. Polar Chromosomes—Challenges of a Risky Path. Cells. 11:1531. doi:10.3390/cells11091531.

53. Wang, P.J., and T.C. Huffaker. 1997. Stu2p: A microtubule-binding protein that is an essential component of the yeast spindle pole body. J Cell Biol. 139:1271–1280. doi:10.1083/jcb.139.5.1271.

54. Zahm, J.A., M.G. Stewart, J.S. Carrier, S.C. Harrison, and M.P. Miller. 2021. Structural basis of Stu2 recruitment to yeast kinetochores. Elife. 10:e65389. doi:10.7554/eLife.65389.

