## Supplemental Figure S1, Table S1, Table S2, Table S3 for "Accurate chromosome segregation is more dependent on kinetochore-bound Stu2 in meiosis than mitosis"

**Supplemental Files**

**Figure S1. Stu2 kinetochore depletion reduces cell viability.** (A) Ten-fold serial dilutions of the indicated strains were spotted onto YPD plates or YPD plates with 1μg/ml of rapamycin. The plates were incubated at 23°C for 36–40 hours before imaging.

**Table S1. Budding yeast strains used in this study.** Strains are derivatives of W303.

**Table S2. Reagents used in this study.**

**Table S3. Plasmids and Primers used in this study.**

**Figure S1.**

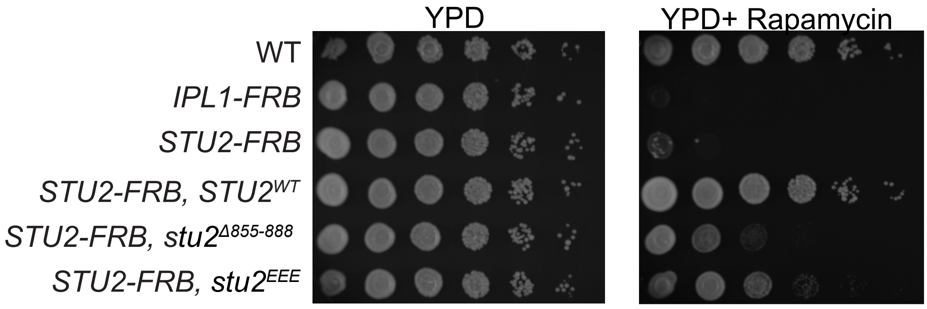

YPD
YPD+ Rapamycin
WT
IPL1-FRB
STU2-FRB
STU2-FRB, STU2WT
STU2-FRB, stu2Δ855-888
STU2-FRB, stu2EEE

### Supplemental table S1. Budding yeast strains used in this study

| Strain | Genotype |
|---|---|
| LY3617 | *MAT**a**/α, tor1-1:HIS3/tor1-1, fpr1::loxP-LEU2-loxP/fpr1::natMX4, RPL13A-2XFKBP12:loxP/RPL13A-2XFKBP12:TRP1, SPC42-mCherry:hphNT1/+, $_{TUB1pr}$GFP-TUB1:URA3/+, ZIP1-GFP/+* |
| LY10982 | *MAT**a**/α, tor1-1:HIS3/tor1-1:HIS3, fpr1::NATMX4/fpr1::NATMX4, RPL13A-2XFKBP12:TRP1/RPL13A-2XFKBP12:TRP1,STU2-FRB:kanMX6/STU2-FRB:kanMX6, SPC42-mCherry:hph, $_{TUB1pr}$GFP-TUB1:URA3/$_{TUB1pr}$GFP-TUB1:URA3* |
| LY10302 | *MAT**a**/α, tor1-1:HIS3/tor1-1:HIS3, fpr1::NATMX4/fpr1::NATMX4, RPL13A-2XFKBP12:TRP1/RPL13A-2XFKBP12:TRP1, $_{TUB1pr}$GFP-TUB1:URA3/$_{TUB1pr}$GFP-TUB1:URA3, SPC42-mCherry:hph, STU2-FRB:kanMX6/STU2-FRB:kanMX6, STU2-3V5:LEU/STU2-3V5:LEU* |
| LY10303 | *MAT**a**/α, tor1-1::HIS3/tor1-1::HIS3, fpr1::NATMX4/fpr1::NATMX4, RPL13A-2XFKBP12::TRP1/RPL13A-2XFKBP12::TRP1, $_{TUB1pr}$GFP-TUB1:URA3/$_{TUB1pr}$GFP-TUB1:URA33, SPC42-mCherry:hph, Stu2-FRB:kanMX6/Stu2-FRB:kanMX6, stu2(855-888)-3V5:LEU/stu2(855-888)-3V5:LEU* |
| LY10304 | *MAT**a**/α, tor1-1:HIS3/tor1-1:HIS3, fpr1::NATMX4/fpr1::NATMX4, RPL13A-2XFKBP12:TRP1/RPL13A-2XFKBP12:TRP1, $_{TUB1pr}$GFP-TUB1:URA3/$_{TUB1pr}$GFP-TUB1:URA3, SPC42-mCherry:hph<br>STU2-FRB:kanMX6/STU2-FRB:kanMX6, stu2(M876E, I873E, L869E mutations)-3V5: LEU/stu2(M876E, I873E, L869E mutations)-3V5:LEU* |
| LY4671 | *MAT**a**/α, tor1-1:HIS3/tor1-1, fpr1::loxP-LEU2-loxP/fpr1::natMX4, RPL13A-2XFKBP12:TRP1/RPL13A-2XFKBP12:TRP1, mad3::kanMX6/mad3::kanMX6, SPC42-mCherry:hphNT1/+, $_{TUB1pr}$GFP-TUB1:URA3/+, ZIP1-GFP/+* |
| LY10480 | *MAT**a**/α, tor1-1:HIS3/tor1-1:HIS3, fpr1::NATMX4/fpr1::NATMX4,RPL13A-2XFKBP12:TRP1/RPL13A-2XFKBP12:TRP1, STU2-FRB:kanMX6/STU2-FRB:kanMX6, mad3::kanMX/mad3::kanMX, SPC42-mCherry:hph,$_{TUB1pr}$GFP-TUB1:URA3/$_{TUB1pr}$GFP-TUB1:URA3* |
| LY10793 | *MAT**a**/α, tor1-1:HIS3/tor1-1:HIS3, fpr1::NATMX4/fpr1::NATMX4,RPL13A-2XFKBP1::TRP1/RPL13A-2XFKBP12:TRP1, STU2-FRB:kanMX6/STU2-FRB:kanMX6, mad3::kanMX/mad3::kanMX, SPC42-mCherry:hph, $_{TUB1pr}$GFP-TUB1:URA3/$_{TUB1pr}$GFP-TUB1:URA3, STU2-3V5:LEU/STU2-3V5:LEU* |
| LY10794 | *MAT**a**/α, tor1-1:HIS3/tor1-1:HIS3, fpr1::NATMX4/fpr1::NATMX4, RPL13A-2XFKBP12:TRP1/RPL13A-2XFKBP12:TRP1, STU2-FRB:kanMX6/STU2-FRB:kanMX6,mad3::kanMX/mad3::kanMX,SPC42-mCherry:hph,$_{TUB1pr}$GFP-TUB1:URA3/$_{TUB1pr}$GFP-TUB1:URA3, stu2(855-888),3V5:LEU/stu2(855-888)-3V5:LEU* |
| LY10795 | *MAT**a**/α,tor1-1:HIS3/tor1-1:HIS3,fpr1::NATMX4/fpr1::NATMX4,RPL13A-2XFKBP12:TRP1/RPL13A-2XFKBP12:TRP1,STU2-FRB:kanMX6/STU2-FRB:kanMX6,mad3::kanMX/mad3::kanMX,SPC42-mCherry:hph,$_{TUB1pr}$GFP-TUB1:URA3/$_{TUB1pr}$GFP-TUB1:URA3, stu2(3M876E, I873E, L869E mutations)-3V5:LEU/Stu2(M876E, I873E, L869E mutations)-3V5:LEU* |
| LY3992 | *MAT**a**/α, tor1-1:HIS3/tor1-1, fpr1::loxP-LEU2-loxP/fpr1::natMX4, RPL13A-2XFKBP12:loxP/RPL13A-2XFKBP12:TRP1, HTB2-mCherry:HIS3/+, $_{TUB1pr}$GFP-TUB1:URA3/+, ZIP1-GFP/+* |

| | |
|---|---|
| LY10941 | *MAT**a**/α, tor1-1/tor1-1, fpr1::NATMX4/fpr1::NATMX4, RPL13A-2XFKBP12:loxP/RPL13A-2XFKBP12:loxP,* $_{TUB1pr}$*GFP-TUB1:URA3,STU2-FRB:kanMX6/STU2-FRB:kanMX6,STU2-3V5:LEU/STU2-3V5:LEU,HTB2-mCherry:HIS3* |
| LY10940 | *MAT**a**/α,tor1-1/tor1-1,fpr1::loxP-LEU2-loxP/fpr1::loxP-LEU2-loxP,RPL13A-2XFKBP12:loxP/RPL13A-2XFKBP12:TRP1,STU2-FRB:kanMX6/STU2-FRB:kanMX6,*$_{TUB1pr}$*GFP-TUB1:URA3/*$_{TUB1pr}$*GFP-TUB1:URA3,HTB2-mCherry:HIS3/-* |
| LY10942 | *MAT**a**/α,tor1-1/tor1-1,fpr1::NATMX4/fpr1::NATMX4,RPL13A-2XFKBP12:loxP/RPL13A-2XFKBP12:loxP,* $_{TUB1pr}$*GFP-TUB1:URA3,STU2-FRB:kanMX6/STU2-FRB:kanMX6,stu2(855-888)-3V5:LEU/stu2(855-888)-3V5:LEU, HTB2-mCherry:HIS3/-* |
| LY10943 | *MAT**a**/α,tor1-1/tor1-1,fpr1::NATMX4/fpr1::NATMX4,RPL13A-2XFKBP12:loxP/RPL13A-2XFKBP12:loxP,* $_{TUB1pr}$*GFP-TUB1:URA3,STU2-FRB:kanMX6/STU2-FRB:kanMX6,stu2(M876E, I873E, L869E mutations)-3V5:LEU/stu2(M876E, I873E, L869E mutations)-3V5:LEU, HTB2-mCherry:HIS3/-* |
| LY11115 | *MAT **a**,tor1-1, fpr1::NATMX4, RPL13A-2XFKBP12::loxP,cup1pr-LacI-GFP:His3, STU2-FRB:kanMX6 LacO:TRP1 (chromosome IV),* $_{TUB1pr}$*mRuby2-TUB1:URA3,stu2-3V5:LEU* |
| LY11116 | *MAT **a**, tor1-1, fpr1::NATMX4, RPL13A-2XFKBP12:loxP,cup1pr-LacI-GFP:His3, STU2-FRB:kanMX6 LacO:TRP1 (chromosome IV),* $_{TUB1pr}$*mRuby2-TUB1:URA3,stu2(855-888)-3V5:LEU* |
| LY11117 | *MAT α,tor1-1, fpr1::NATMX4, RPL13A-2XFKBP12:loxP,cup1pr-LacI-GFP:His3, STU2-FRB:kanMX6 LacO:TRP1 (chromosome IV),* $_{TUB1pr}$*mRuby2-TUB1:URA3,stu2-3V5:LEU* |
| LY11118 | *MAT α, tor1-1, fpr1::NATMX4, RPL13A-2XFKBP12:loxP,cup1pr-LacI-GFP:His3, STU2-FRB:kanMX6 LacO:TRP1 (chromosome IV),* $_{TUB1pr}$*mRuby2-TUB1:URA3, stu2(855-888)-3V5:LEU* |
| LY11095 | *MAT **a**, tor1-1, fpr1::NATMX4, RPL13A-2XFKBP12:loxP,cup1pr-LacI-GFP:His3, STU2-FRB:kanMX6 LacO:TRP1 (chromosome IV),* $_{TUB1pr}$*mRuby2-TUB1:URA3, stu2(M876E, I873E, L869E mutations)-3V5 :LEU* |
| LY11096 | *MAT α, tor1-1, fpr1::NATMX4, RPL13A-2XFKBP12:loxP,cup1pr-LacI-GFP:His3, STU2-FRB:kanMX6 LacO:TRP1 (chromosome IV),* $_{TUB1pr}$*mRuby2-TUB1:URA3, stu2(M876E, I873E, L869E mutations)-3V5 :LEU* |
| LY11119 | *MAT **a**,tor1-1,fpr1::NATMX4,RPL13A-2XFKBP12:TRP1,STU2-FRB:kanMX6,HTB2-GFP:TRP1 ,stu2-3V5:LEU* |
| LY11120 | *MAT **a**,tor1-1,fpr1::NATMX4,RPL13A-2XFKBP12:TRP1,STU2-FRB:kanMX6,HTB2-GFP:TRP1,<br>stu2(855-888)-3V5:LEU* |
| LY11121 | *MAT **a**,tor1-1,fpr1::NATMX4,RPL13A-2XFKBP12:TRP1,STU2-FRB:kanMX6,HTB2-GFP:TRP1 ,stu2(M876E, I873E, L869E mutations)-3V5:LEU* |
| LY11122 | *MAT α,tor1-1,fpr1::NATMX4,RPL13A-2XFKBP12:TRP1,STU2-FRB:kanMX6, SPC42-RedStar2:NAT, stu2-3V5:LEU* |
| LY11123 | *MAT α,tor1-1,fpr1::NATMX4,RPL13A-2XFKBP12:TRP1,STU2-FRB:kanMX6, SPC42-RedStar2:NAT,stu2(855-888)-3V5:LEU* |
| LY11124 | *MAT α, tor1-1,fpr1::NATMX4,RPL13A-2XFKBP12:TRP1,STU2-FRB:kanMX6, SPC42-RedStar2:NAT,stu2(M876E, I873E, L869E mutations)-3V5:LEU* |
| LY10950 | *MAT**a**/α, tor1-1:HIS3/tor1-1:HIS3,fpr1::NATMX4/fpr1::NATMX4, RPL13A-2XFKBP12:loxp/RPL13A-2XFKBP12:loxp,*$_{TUB1pr}$*GFP-TUB1:URA3/*$_{TUB1pr}$*GFP-TUB1:URA3,STU2-FRB:kanMX6/STU2-FRB:kanMX6,IPL11-FRB:kanMX6/IPL1-FRB:kanMX6,SPC42-mCherry:hph* |

| | |
|---|---|
| LY11004 | *MAT α, tor1-1:HIS3, fpr1::NATMX4, RPL13A-2XFKBP12:loxp, $_{TUB1pr}$GFP-TUB1:URA3, STU2-FRB:kanMX6, IPL1-FRB:kanMX6, stu2-3V5:LEU* |
| LY11005 | *MAT **a**, tor1-1:HIS3, fpr1::NATMX4, RPL13A-2XFKBP12:loxp, $_{TUB1pr}$GFP-TUB1:URA3, STU2-FRB:kanMX6, IPL1-FRB:kanMX6, stu2-3V5:Leu, SPC42-mCherry:hph* |
| LY11006 | *MAT α, tor1-1:HIS3, fpr1::NATMX4, RPL13A-2XFKBP12:loxp, $_{TUB1pr}$GFP-TUB1:URA3, STU2-FRB:kanMX6, IPL1-FRB:kanMX6, stu2(855-888)-3V5:LEU* |
| LY11007 | *MAT **a**, tor1-1:HIS3, fpr1::NATMX4, RPL13A-2XFKBP12:loxp, $_{TUB1pr}$GFP-TUB1:URA3, STU2-FRB:kanMX6, IPL1-FRB:kanMX6,stu2(855-888)-3V5:LEU, SPC42-mCherry:hph* |
| LY11008 | *MAT α, tor1-1:HIS3, fpr1::NATMX4, RPL13A-2XFKBP12:loxp, $_{TUB1pr}$GFP-TUB1:URA3, STU2-FRB:kanMX6, IPL1-FRB:kanMX6, stu2(M876E, I873E, L869E mutations)-3V5:LEU* |
| LY11009 | *MAT **a**, tor1-1:HIS3, fpr1::NATMX4, RPL13A-2XFKBP12:loxp, $_{TUB1pr}$GFP-TUB1:URA3, STU2-FRB:kanMX6, IPL1-FRB:kanMX6, stu2(M876E, I873E, L869E mutations)-3V5:LEU, SPC42-mCherry:hph* |
| LY11027 | *MAT α, tor1-1:HIS3, fpr1::NATMX4, RPL13A-2XFKBP12:loxp, $_{TUB1pr}$GFP-TUB1:URA3, STU2-FRB:kanMX6, IPL1-FRB:kanMX6, Mad3::hph (3HA),SPC42-mCherry:hph* |
| LY11028 | *MAT **a**, tor1-1:HIS3, fpr1::NATMX4, RPL13A-2XFKBP12:loxp, $_{TUB1pr}$GFP-TUB1:URA3, STU2-FRB:kanMX6, IPL1-FRB:kanMX6, Mad3::hph (3HA)* |
| LY11043 | *MAT α, tor1-1:HIS3, fpr1::NATMX4, RPL13A-2XFKBP12:loxp, $_{TUB1pr}$GFP-TUB1:URA3, STU2-FRB:kanMX6, IPL1-FRB:kanMX6, Mad3::hph (3HA),SPC42-mCherry:hph, stu2(855-888)-3V5:LEU* |
| LY11044 | *MAT α, tor1-1:HIS3, fpr1::NATMX4, RPL13A-2XFKBP12:loxp, $_{TUB1pr}$GFP-TUB1:URA3, STU2-FRB:kanMX6, IPL1-FRB:kanMX6, Mad3::hph (3HA),SPC42-mCherry:hph, stu2(M876E, I873E, L869E mutations)-3V5:LEU* |
| LY11045 | *MAT **a**, tor1-1:HIS3, fpr1::NATMX4, RPL13A-2XFKBP12:loxp, $_{TUB1pr}$GFP-TUB1:URA3, STU2-FRB:kanMX6, IPL1-FRB:kanMX6, Mad3::hph (3HA),stu2(855-888)-3V5:LEU* |
| LY11046 | *MAT **a**, tor1-1:HIS3, fpr1::NATMX4, RPL13A-2XFKBP12:loxp, $_{TUB1pr}$GFP-TUB1:URA3, STU2-FRB:kanMX6, IPL1-FRB:kanMX6, Mad3::hph (3HA),stu2(M876E, I873E, L869E mutations)-3V5:LEU* |
| LY11151 | *MAT **a**, tor1-1:HIS3, fpr1::NATMX4, RPL13A-2XFKBP12:loxp, $_{TUB1pr}$GFP-TUB1:URA3, STU2-FRB:kanMX6, IPL1-FRB:kanMX6, HTB2-mCherry:Hph, stu2-3V5:LEU* |
| LY11152 | *MAT **a**, tor1-1:HIS3, fpr1::NATMX4, RPL13A-2XFKBP12:loxp, $_{TUB1pr}$GFP-TUB1:URA3, STU2-FRB:kanMX6, IPL1-FRB:kanMX6, HTB2-mCherry:Hph, stu2(855-888)-3V5:LEU* |
| LY11153 | *MAT **a**, tor1-1:HIS3, fpr1::NATMX4, RPL13A-2XFKBP12:loxp, $_{TUB1pr}$GFP-TUB1:URA3, STU2-FRB:kanMX6, IPL1-FRB:kanMX6, HTB2-mCherry:Hph, stu2(M876E, I873E, L869E mutations)-3V5:LEU* |
| LY11173 | *MAT **a**, tor1-1, fpr1::NATMX4, RPL13A-2XFKBP12:loxP, STU2-FRB:kanMX6, IPL1-FRB:kanMX6, LacO:TRP1 (chromosome IV), $_{TUB1pr}$mRuby2-TUB1:URA3, stu2-3V5:LEU* |
| LY11174 | *MAT α, tor1-1, fpr1::LoxP-LEU2-loxP, RPL13A-2XFKBP12:loxP, STU2-FRB:kanMX6, IPL1-FRB:kanMX6,cup1pr-LacI-GFP:His3, LacO:TRP1 (chromosome IV), stu2-3V5:LEU* |
| LY11175 | *MAT α, tor1-1 fpr1::Nat RPL13A-2XFKBP12:loxP cup1pr-LacI-GFP:His3 STU2-FRB:kanMX6 LacO:TRP1 (chromosome IV) IPL1-FRB:kanMX6, $_{TUB1pr}$mRuby2-TUB1:URA3* |
| LY11176 | *MAT **a**, tor1-1 fpr1::Nat RPL13A-2XFKBP12:loxP, STU2-FRB:kanMX6, LacO:TRP1 (chromosome IV), IPL1-FRB:kanMX6* |
| LY11177 | *MAT **a**, tor1-1: fpr1::NATMX4 RPL13A-2XFKBP12:loxP, $_{TUB1pr}$GFP-TUB1:URA3,STU2-FRB:kanMX6, stu2(855-888)-3V5:LEU, HTB2-mCherry:His* |
| LY11178 | *MAT **a**, tor1-1: fpr1::NATMX4 RPL13A-2XFKBP12:loxP, $_{TUB1pr}$GFP-TUB1:URA3 STU2-FRB:kanMX6, stu2(M876E, I873E, L869E mutations)-3V5:LEU, HTB2-mCherry:His* |

| | |
|---|---|
| LY11181 | MAT α, tor1-1 fpr1::NAT, RPL13A-2XFKBP12:loxP, cup1pr-LacI-GFP:His3, STU2-FRB:kanMX6, LacO:TRP1 (chromosome IV), IPL1-FRB:kanMX6, $_{TUB1pr}$mRuby2-TUB1:URA3, stu2(M876E, I873E, L869E mutations)-3V5:LEU |
| LY11182 | MAT **a**, tor1-1 fpr1::Nat, RPL13A-2XFKBP12:loxP, STU2-FRB:kanMX6, LacO:TRP1 (chromosome IV), IPL1-FRB:kanMX6, stu2(M876E, I873E, L869E mutations)-3V5:LEU |
| LY11211 | MAT α, tor1-1 fpr1::Nat, RPL13A-2XFKBP12:loxP, cup1pr-LacI-GFP:His3, STU2-FRB:kanMX6, LacO:TRP1 (chromosome IV), IPL1-FRB:kanMX6, $_{TUB1pr}$mRuby-TUB1:URA3, stu2(855-888)-3V5:LEU |
| LY11212 | MAT **a**, tor1-1 fpr1::Nat, RPL13A-2XFKBP12:loxP, stu2-FRB:kanMX6, LacO:TRP1 (chromosome IV), IPL1-FRB:kanMX6, stu2(855-888)-3V5:LEU |
| LY5725 | MAT**a**/α, tor1-1:HIS3/tor1-1, fpr1::NATMX4/fpr1::loxP-LEU2-LoxP, RPL13A-2XFKBP12:TRP1/RPL13A-2XFKBP12:loxP , $_{TUB1pr}$GFP-TUB1:URA3/+ SPC42-mCherry:hph/+ IPL1-FRB:kanMX6/IPL1-FRB:kanMX6, Zip1-GFP/ |
| LY5703 | MAT **a**, tor1-1, fpr1::LoxP-LEU2-loxP, RPL13A-2XFKBP12:loxP, IPL1-FRB:kanMX6, LacO:TRP1, cup1pr-LacI-GFP:His3 |
| LY11246 | |
| LY5149 | MAT α, tor1-1:HIS3 fpr1::NATMX4, RPL13A-2XFKBP12::TRP1, $_{TUB1pr}$GFP-TUB1:URA3, SPC42-mCherry:hph, IPL1-FRB:kanMX6 |
| LY6039 | MAT **a**, tor1-1, fpr1::LoxP-LEU2-loxP, RPL13A-2XFKBP12:,:loxP, SPC42-RedStar2:NAT, HTB2-GFP:TRP1, IPL1-FRB:kanMX6 |
| LY5522 | MAT **a**, tor1-1, fpr1::NATMX4, RPL13A-2XFKBP12:loxP, $_{TUB1pr}$GFP-TUB1:URA3, SPC42-mCherry:hph |
| LY8229 | MAT α, tor1-1:HIS3, fpr1::NATMX4, RPL13A-2XFKBP12:TRP1, STU2-FRB:kanMX6, SPC42-mCherry:hph, $_{TUB1pr}$GFP-TUB1:URA3 |
| LY5165 | MAT**a**/α, tor1-1:HIS3/tor1-1, fpr1::loxP-LEU2-loxP/fpr1::natMX4, RPL13A-2XFKBP12:loxP/RPL13A-2XFKBP12:TRP1, IPL1-FRB:kanMX6/IPL1-FRB:kanMX6, HTB2-mCherry:HIS3/+, $_{TUB1pr}$GFP-TUB1:URA3/+, ZIP1-GFP/+ |
| LY5166 | MAT**a**/α, tor1-1:HIS3/tor1-1, fpr1::loxP-LEU2-loxP/fpr1::natMX4, RPL13A-2XFKBP12:loxP/RPL13A-2XFKBP12:TRP1, IPL1-FRB:kanMX6/IPL1-FRB:kanMX6, SPC42-mCherry:hphNT1/+,$_{TUB1pr}$GFP-TUB1:URA3/+, ZIP1-GFP/+ |
| LY4495 | MAT**a**/α, tor1-1/tor1-1,fpr1::LoxP-LEU2-loxP/fpr1::NAT, RPL13A-2XFKBP12::loxP/RPL13A-2XFKBP12::TRP1, HTB2-GFP:TRP1/+SPC42-RedStar2:NAT/+ |

**Supplemental table S2. Reagents used in this study**

| Reagent | Source | Identifier |
|---|---|---|
| Yeast extract | Gibco | Ref#212720 Lot#2943970 |
| Dextrose (D-Glucose) Anhydrous | Fisher Scientific | CAS#50-99-7 Lot#225156 |
| Peptone | Gibco | Ref# 211677 Lot#5141151 |
| L-Tryptophan | Sigma-Aldrich | CAS#73-22-3 Lot#SLBM3572V |
| Adenine hemisulfate salt | Sigma-Aldrich | CAS# 321-30-2 Lot#WXBD0156V |
| Bacto-agar | Sunrise Science | Cat#1910-5KG Lot#26B3673 |
| Potassium acetate | Fisher Scientific | Lot#219374 |
| Synthetic complete mixture drop-out: Complete | Sunrise Science | Cat#1300-030 Lot#23M3191 |
| Yeast nitrogen base without amino acids | Fisher Scientific | Ref#291920 Lot#9148845 |
| NaCl | EMD Chemicals | CAS#7647-14-5 Lot#49012905 |
| Tryptone (Casein Peptone) | Sunrise Science | Cat#1914-1KG Lot#5K0771 |
| Ampicillin sodium salt | Sigma-aldrich | Cat#A9518-5G |
| Rapamycin | Thermo Fisher Scientific | Cat# AAJ62473MF |
| Alpha factor | Zymo Research | Cat#Y1001 |
| Concanavalin A | Sigma-Aldrich | CAS#11028-71-0 Lot#0000266073 |
| Nourseothricin Sulfate (Clonat) | GBiosciences | Cat#RC-187 Lot#221801 |
| G418 sulphate | Fisher Bioreagents | Cat#BP673-5 |
| Hygromycin B Cornig | Invitrogen | Cat#10687010 |
| Dimethyl sulfoxide (DMSO) | Sigma-aldrich | Cat#5879-500ML |
| Ethanol | KOPTEC | CAS#64-17-5 |
| Glycerol | Invitrogen | Cat# 15514029 |
| Phenol/chloroform/isoamyl alcohol | Fisher Bioreagents | Cat#BP1752I-400 |
| single-stranded salmon sperm DNA | Sigma-aldrich | Cat#D1626-5G |
| Lithium acetate dihydrate | Sigma-aldrich | Cat#L6883-1KG |
| Polyethylene glycol | Spectrum chemical MFG corp | Cat#25322-68-3 |

**Supplemental Table S3. Plasmids and Primers used in this study**

| | | Source | |
|---|---|---|---|
| pLB574 | $_{STU2pr}$*STU2-3V5:LEU2* | 1 | Gift of M. Miller |
| pLB576 | $_{STU2pr}$*stu2(Δ855-888)-3V5:LEU2* | 1 | Gift of M. Miller |
| pLB577 | $_{STU2pr}$*stu2(M876E I873E L869E)-3V5:LEU2* | 1 | Gift of M. Miller |
| pLB113 | $_{CUP1pr}$*LacI-GFP:HIS3* | | Gift of A.W. Murray |
| pSLB21 | LacO:Trp1 | 2 | |
| LO2831 | 5'ATATTTTATTGAGTTTATGTTATGGGGAGGCTA<br>CCCTTTA<br>gaattcgagctcgtttaaac-3' | This study | To tag stu2 with FRB |
| LO2832 | 5'ATATTTTATTGAGTTTATGTTATGGGGAGGCTA<br>CCCTTTAgaattcgagctcgtttaaac-3' | This study | To tag stu2 with FRB |
| LO2833 | 5'-GCAGTGTCTCCGACGATTTG-3' | This study | stu2-Ctertag-checkF |
| LO2834 | 5'-CCAGTCGCATGTTGTTACCC-3' | This study | stu2-Ctertag-checkR |
| LO3607 | 5'GTTGTTAGAGAACTAGTGGGTGG-3' | This study | To check that Stu2 is integrated at Leu2. |
| LO3611 | 5'-GTCGCTATTGTTGCCCGTAACCT-3' | This study | To check Stu2 is integrated at Leu2. |
| LO1751 | 5'GATACTAAGAAACAAGCCCTTTTGGGAAAATA<br>AGCGGTTAcggatccccgggttaattaa-3' | This study | To tag Ipl1 with FRB |
| LO1752 | 5'AATAGTGCCCTTCAAACGATTCTGTCATACTT<br>TAATTCTAgaattcgagctcgtttaaac-3' | This study | To tag Ipl1 with FRB |
| LO1753 | 5'-CTGCGAATGCTCGTCTTGAG-3' | This study | To check FRB tag (or C terminal tag) on Ipl1 |
| LO1754 | 5'-CGTCCTGGCGTTTGAACTAC-3' | This study | To check FRB tag (or C terminal tag) on Ipl1 |


1. Zahm JA, Stewart MG, Carrier JS, Harrison SC, Miller MP. Structural basis of Stu2 recruitment to yeast kinetochores. Elife. 2021 Feb 16;10:e65389. doi: 10.7554/eLife.65389. PMID: 33591274; PMCID: PMC7909949.
2. Straight AF, Belmont AS, Robinett CC, Murray AW. GFP tagging of budding yeast chromosomes reveals that protein-protein interactions can mediate sister chromatid cohesion. Current Biology : CB. 1996;6(12):1599-608. Epub 1996/12/01. doi: 10.1016/s0960 9822(02)70783-5. PubMed PMID: 8994824.
